# Decoding mEos4b Day-Long Maturation and Engineering Fast Maturing Variants

**DOI:** 10.1101/2024.09.26.615204

**Authors:** Arijit Maity, Oleksandr Glushonkov, Isabel Ayala, Pascale Tacnet, Jip Wulffelé, Philippe Frachet, Bernhard Brutscher, Dominique Bourgeois, Virgile Adam

## Abstract

The maturation speed of fluorescent proteins is a crucial parameter that influences cellular brightness, effective labeling efficiency and temporal resolution in fluorescence microscopy. Green-to-red photoconvertible fluorescent proteins (PCFPs) used in pulse-chase experiments and super-resolution techniques such as Photoactivated Localization Microscopy (PALM), single-particle-tracking PALM (sptPALM) and Minimal Fluorescence Photon Fluxes Microscopy (MINFLUX) may be hampered by slow maturation. We systematically characterized the maturation speed of mEos-derived PCFPs in *E. coli* and found that, in contrast to pcStar and mEosEM, several variants such as mEos2, mEos3.1, mEos3.2 and mEos4b mature extremely slowly. Strikingly, the oxidation step in those PCFPs is fast and not rate-limiting. Through a rational mutagenesis approach, we developed a strategy to reduce the day-long maturation time of mEos4b by nearly two orders of magnitude without significantly impacting its molecular brightness and photophysical performance under single-molecule imaging conditions.

## Introduction

Fluorescent proteins have revolutionized the field of cell biology, serving as indispensable tools for visualizing and tracking cellular processes. Their ability to act as non-invasive markers has significantly contributed to our understanding of complex biological phenomena. However, a persistent challenge for the utility of fluorescent proteins is the potentially slow maturation of their chromophore. The rate of chromophore maturation is a critical factor that affects various aspects of cellular imaging, including apparent brightness, effective labelling efficiency and temporal resolution. Slow maturation may also result in artefacts in FRET measurements^1, 2^ and the toxic accumulation of hydrogen peroxide in cells.^3^ Photoconvertible fluorescent proteins (PCFPs) are commonly employed in super-resolution microscopy techniques such as Photo-Activated Localization Microscopy (PALM), single-particle-tracking PALM (sptPALM) and more recently MINFLUX.^4^ Slow maturation can be a notable drawback since it may hamper the successful visualization and analysis of cellular structures at enhanced resolutions, notably in live cells. In this work, we present a comprehensive investigation of the maturation speed of PCFPs from the mEos family and propose strategies to considerably reduce the maturation time of the monomeric, bright and fixation-resistant mEos4b variant.^5^

The mechanism by which the chromophore of fluorescent proteins matures has been widely studied, but remains incompletely understood. Following protein folding, the carbonyl function of the first amino acid in the chromophoric triad (S65 for EGFP but variable among FPs) and the amide function of the third amino acid (G67, strictly conserved) come in proximity, as a result of the prototypical kinked central α-helix of the FP scaffold.^6^ In this orientation, nucleophilic attack of the nitrogen from G67^7, 8^ on the carbonyl of S65 (in EGFP) leads to the formation of a covalent bond in a process called cyclisation. This cyclisation step forms an initial intermediate with an imidazolidine ring. The subsequent reactions involve the formation of double bonds through an oxidation step and a dehydration step. Depending on the order of these two steps, two competing models have been proposed (Figure 1). According to the first model (Figure 1, cyan arrows) the dehydration step precedes the oxidation step.^9-17^ Elimination of a hydroxyl group and one proton result in the formation of a first double bond in an imidazolinone ring. Molecular oxygen attack on the chromophore then results in a (hydro)peroxy intermediate, leading to the elimination of hydrogen peroxide and the formation of a second double bond establishing conjugation between the hydroxybenzylidene and imidazolinone groups.

**Figure 1.**
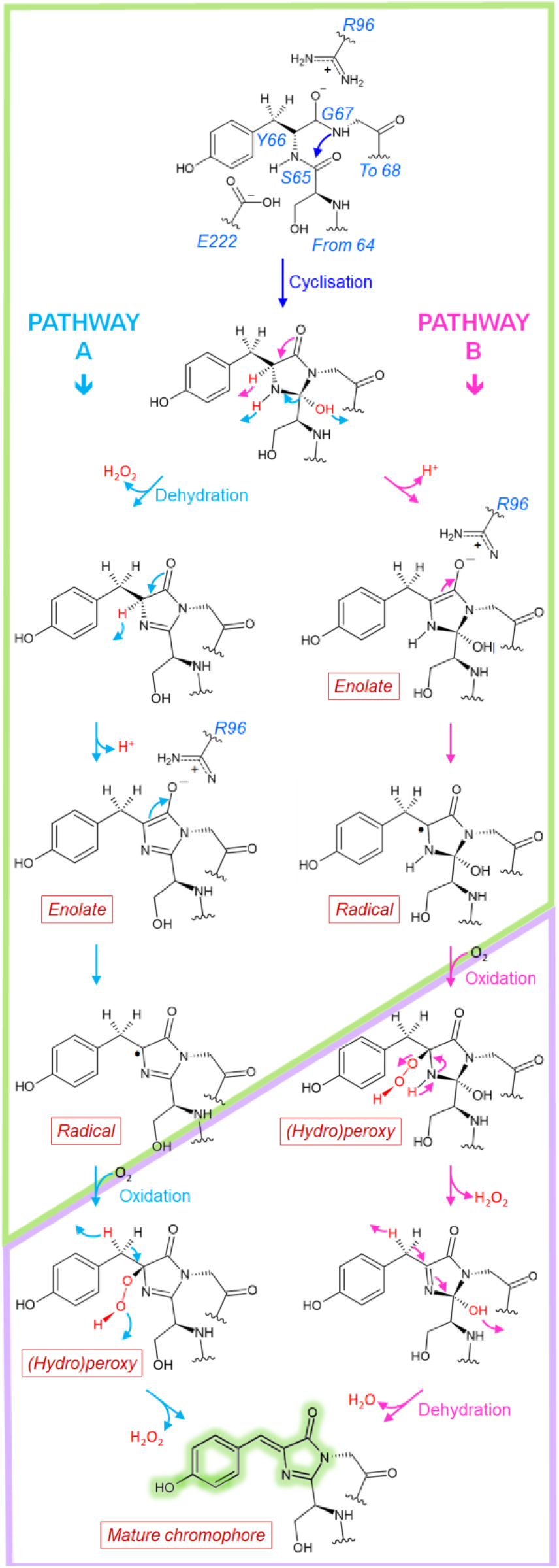
Chemical reaction steps of maturation from the pre cyclised to the mature chromophore in EGFP. Following the cyclisation step (blue), two possible pathways have been proposed. In the first pathway (cyan), a dehydration step followed by an oxidation step involving molecular oxygen, results in the elimination of two hydrogen atoms as hydrogen peroxide. In the alternative pathway (pink), these steps are reversed. Both pathways result in the formation of double bonds, allowing the establishment of a conjugated system in the mature chromophore. Leaving groups are shown in red. The oxygen-independent steps are framed in green while the oxygen-dependent steps are framed in purple.

According to the second model (Figure 1, pink arrows), the oxidation step precedes the dehydration step.^18-21^ Molecular oxygen attack results in a (hydro)peroxy intermediate, and following the elimination of hydrogen peroxide, a new hydroxyl imidazolinone intermediate is formed. Dehydration then leads to the mature chromophore.

The possibility of both pathways acting in parallel has also been discussed.^22^ Regardless of the pathway, these maturation steps are facilitated by the chromophore’s environment, notably the strictly conserved amino acids E222 (numbered in EGFP, E212 in mEos4b) and R96 (numbered in EGFP, R91 in mEos4b). E222 functions as a general base, facilitating deprotonation of the chromophore G67 or abstracting a proton from the α-carbon of Y66, possibly mediated via a water molecule.^7, 23, 24^ Symmetrically, R96 promotes electrophilic catalysis through its interaction with the carbonyl group of G67, notably by stabilizing the enolate intermediate, key to chromophore cyclisation.^8, 10, 13, 25-^

Mutation of these residues is not well tolerated and, for example, the mutation of E222 to a glutamine slows down the maturation process at physiological pH and accelerates it at very basic pH.^23^ The exchange of R96 with a different amino acid (except lysine) was reported to drastically slow down maturation kinetics^8, 26^ up to several months.^6, 23^

Furthermore, the precise positioning and interaction patterns of E222 and R96 within the FP scaffold may affect the maturation rate. To date, mechanistic studies on FP maturation have only been conducted on hydrozoan GFP variants. The emerging picture is that the rate-limiting step in maturation is the oxidation step,^9, 11, 12, 19, 28^ whereas cyclization is a fast process.^9^ However, whether this scenario also applies to FPs of anthozoan origin remains unknown.

Recent advancements have seen the emergence of FP variants with faster maturing chromophores, such as Venus,^28^ DsRed-express^29^ and DsRed-express2,^30^ mScarlet3,^31^ Kohinoor2.0^32^ and mStayGold^33^ compared to their predecessors. Since a general rationale for the improved maturation kinetics is missing, the mutagenesis strategy employed for these FPs cannot be translated directly to improve the maturation speed of other FPs. It is also important to note that maturation rates reported in the literature for single FPs (e.g. mNeonGreen, Clover, mStayGold, mBaojin or Ds-Red express variants)^29, 33-36^ vary significantly. The exact definition of maturation kinetics also differs between studies, and the techniques employed to measure it are diverse (as it is the case for other parameters such as quantum yield or extinction coefficient,^37^) leading to discrepancies in the reported values. For example the re-folding rate following FP denaturation is sometimes (incorrectly) reported as maturation rate.^38^ Consequently, data available in repositories such as FPbase^39^ may not always be reliable. Balleza *et al*. have highlighted the need for a rigorous study of maturation rates in FPs,^36^ but a systematic investigation of PCFP maturation rates has not been carried out until now.

When studying mEos4b by Nuclear Magnetic Resonance (NMR) spectroscopy in the frame of our recent work^,40^ we discovered that this PCFP exhibits exceptionally slow maturation. As this unfavourable property has not been reported previously in the literature, we undertook a detailed investigation of maturation kinetics in proteins of the mEos family. To this aim, we devised a well-defined methodology to measure the apparent maturation rate (from folding to appearance of fluorescence signal, i.e. the parameter of practical importance for imaging applications) of PCFPs. Moreover, we separately measured the rate of the oxidation (and possibly post-oxidation) step.

Our measurements revealed that the two most recent members of the mEos family: pcStar^41^ and mEosEM,^42^ exhibit significantly faster maturation than older variants. To better understand the rationale behind these differences, we identified the key mutations involved. With the help of crystallographic structures, AlphaFold calculations^43^ and MD simulations^,44^ we conducted mutagenesis studies on mEos4b. Among the variants generated, two double mutants named mEos4Fast1 and mEos4Fast2 exhibited markedly accelerated maturation kinetics compared to their parent while retaining other biochemical and photophysical properties. Finally, to validate the practical utility of our findings, we applied the engineered variants for cellular imaging in both prokaryotic and eukaryotic cells.

## Results

### NMR Insights and Comparative Maturation Kinetics

During our recent study of mEos4b photophysics by multidimensional NMR spectroscopy,^40^ we observed that maturation of this PCFP was exceptionally long. NMR ^1^H-^15^N spectra of mEos4b, acquired immediately after protein production and purification, and up to 36 h later while remaining in the NMR magnet at 35°C, revealed a progressive disappearance of NMR peaks corresponding to a well-folded species of mEos4b, tentatively assigned to proteins harbouring an incompletely matured chromophore (Figure 2A-C). Concomitantly, the intensity of the major peak species, corresponding to the green fluorescent state, increased during the kinetic experiment. Comparison of the UV-Vis absorption spectra of a freshly purified sample recorded before and immediately after the NMR experiment (after 36 h) showed a significantly increased absorption band at 504 nm. This provides evidence for incomplete chromophore maturation at the start of NMR data collection (Figure 2D).

**Figure 2.**
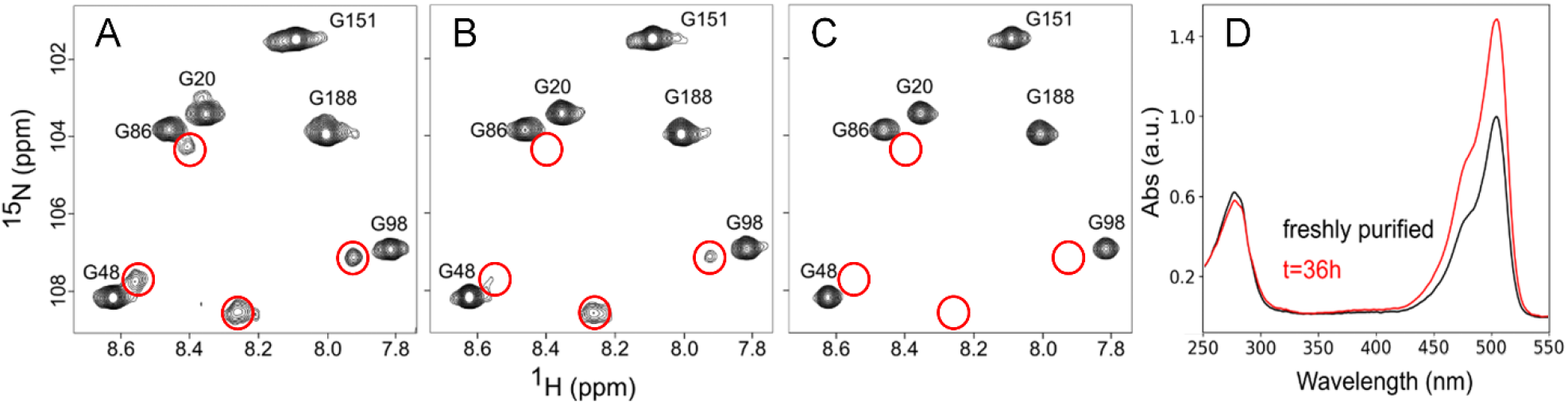
Slow chromophore maturation kinetics observed in NMR and UV-Vis spectroscopy at 35°C. (A-C) Small region of NMR ^1^H-^15^N spectra of mEos4b acquired (A) just after protein production and purification, and after (B) 8 h and (C) 36 h in the NMR magnet. NMR peaks disappearing during maturation of the chromophore are highlighted by red circles. The visible green state peaks are annotated by their residue number and type. (D) Comparison of UV-Vis spectra of freshly purified (black) sample and after 36 h (red) in the NMR magnet.

The evolution of the NMR signal intensity thus provided a measure of apparent maturation kinetics. Fitting the data to a mono-exponential kinetic model yielded a time constant of about 1200 min (Figure S1), that is more than an order of magnitude slower than previously reported values for members of the mEos family, namely mEosFP, mEos2, mEos3.1 and mEos3.2^.38^ We noticed, however, that these literature values essentially reported refolding times of the proteins following chemical denaturation^.38^ We therefore measured the full maturation kinetics of mEos4b and other members of the mEos family with an *in vitro* fluorescence-based assay (see methods).

At 25°C, the fluorescence build-up of mEos4b pointed to an extremely slow maturation compared to the more recent pcStar and mEosEM (Figure 3A), in line with our NMR data. The fluorescence did not reach a plateau even after 5000 min, suggesting the maturation was still incomplete. Furthermore, the fluorescence curve clearly indicated the presence of at least two populations that matured at different rates. The relatively fast maturing population could be fitted with a mono-exponential build-up with a time constant of 1816 ± 45 min. The data also revealed that in spite of very similar sequences (Figure 3B), a significantly slower maturation rate for mEos4b was observed as compared to its ancestors mEos2, mEos3.1 and mEos3.2 (Figure 4, Figure S2). However, all these variants showed maturation times exceeding 11 hours. Only the recent mEos variants, pcStar and mEosEM, exhibited significantly faster maturation with time constants of the order of 30 minutes.

**Figure 3.**
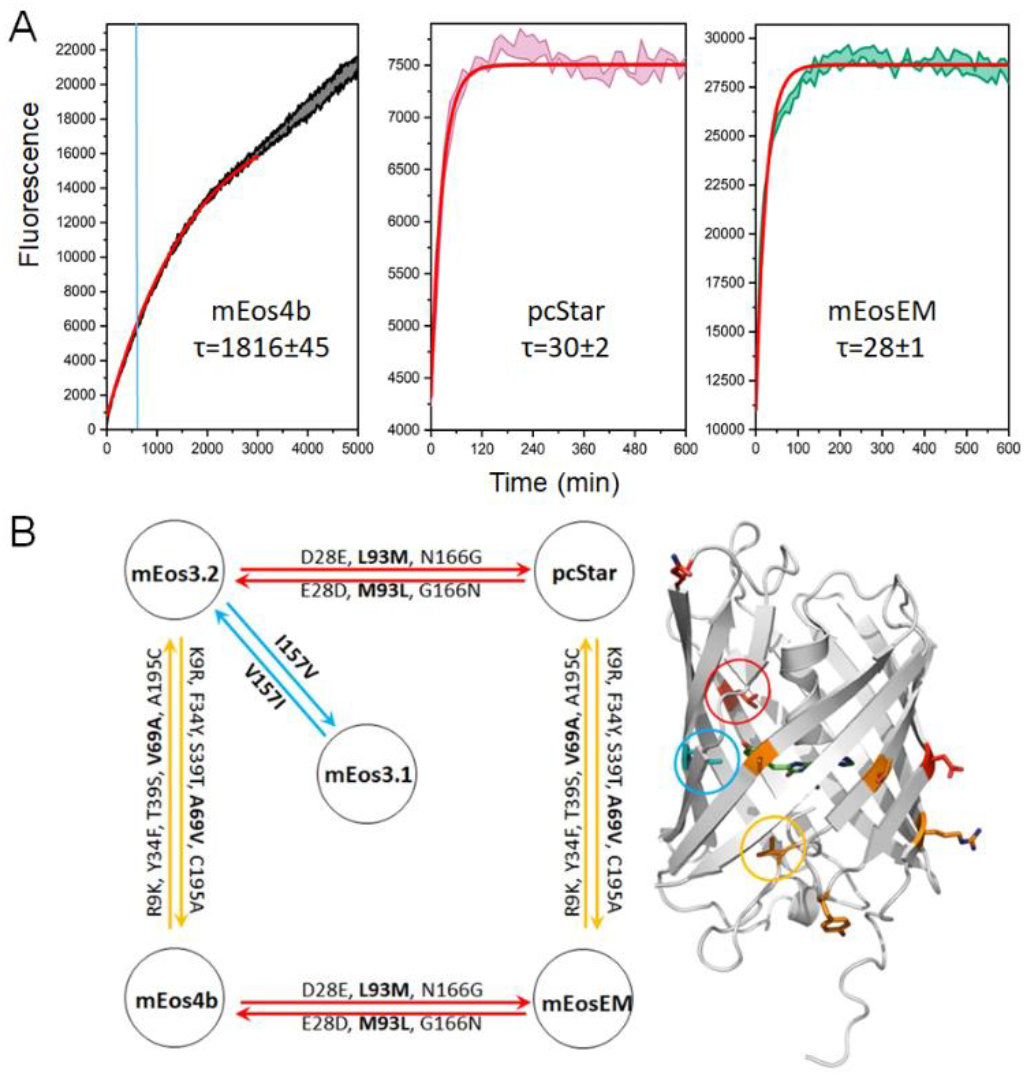
(A) Apparent maturation rates measured by fluorescence increase in bacteria of mEos4b and the two latest mEos derivatives pcStar and mEosEM. All fluorescence measurements were performed at 25°C. For mEos4b the total acquisition time was longer than represented and the vertical blue line represents the time window that was used for pcStar and mEosEM. (B) Mutation map of mEos4b with its closest relatives, namely mEos3.1, mEos3.2, pcStar and mEosEM. All mutations are mapped on the crystal structure of mEos4b, with the mutation sites inside the barrel circled and shown in bold.

**Figure 4.**
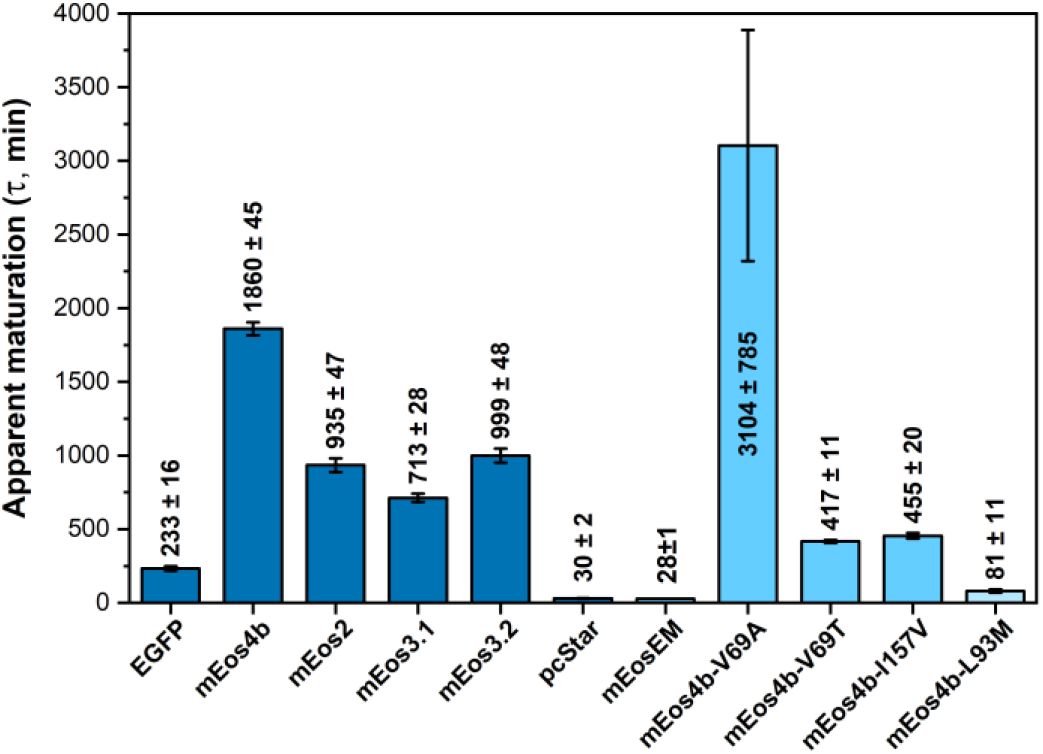
Time constants for the fast phase of apparent maturation kinetics (folding + cyclization + dehydration + oxidation) for EGFP (control) and mEos4b and its ancestors (dark blue) as compared to single mEos4b variants (light blue). Error bars are standard deviations of time constants extracted from three separate measurements.

### Position 93 is key for maturation

In light of the several orders of magnitude differences in maturation rates observed among various mEos relatives, we conducted a detailed comparison of their primary sequences. Figure 3B shows the sequence relationship between mEos4b, mEos3.1, mEos3.2, pcStar and mEosEM. When compared to mEos4b and mEosEM, most of the differences in the other three proteins are located outside the β-barrel. While long range effects have been previously shown to affect properties of fluorescent proteins^,21, 29^ they are difficult to predict. Therefore, we primarily focussed on residues with side chains pointing towards the interior of the β-barrel, as depicted in Figure 3B, specifically positions 69, 93, and 157. As illustrated in the same figure, the only difference between mEos3.1 and mEos3.2 occurs at position 157, which is a valine in mEos3.1 and an isoleucine in mEos3.2. Yet, this single substitution from valine to isoleucine increases the maturation time of mEos3.2 by ∼40% (Figure 4). Similarly, we hypothesized that the L93M mutation, which distinguishes mEos3.2 from pcStar and mEos4b from mEosEM, could be pivotal in the significant reduction in maturation time observed for pcStar and mEosEM. Additionally, we speculated that the A69V mutation could also cause near doubling of maturation time in mEos4b as compared to mEos3.2.

To test these hypotheses, we generated the I157V, L93M, V69A substitutions in mEos4b and measured their maturation kinetics. Additionally, we generated the V69T variant to test a charged residue that we previously identified as important to modify the microenvironment of the chromophore.^45^ Our measurements showed that, while the mutation V69A further slowed down the maturation kinetics of mEos4b, the 3 other single-point mutants improved it. Despite this, the V69T and I157V mutations were still unsatisfactory with maturation time constants exceeding 400 min. In contrast, the L93M mutant showed a drastically decreased apparent maturation time of approximately 80 min, a more than 20-fold improvement relative to the mEos4b parent (Figure 4, Figure S3A) and better than any other L93 substitutions we have tested (Table S1).

Despite this improvement, the maturation rate of mEos4b-L93M remained approximately three-fold slower compared to that of pcStar or mEosEM (Figure 4), both of which also have a methionine at position 93 (Figure 3B). Hence, we set out to find other mutations that would further enhance maturation rates without compromising other photophysical properties, particularly in the photoconverted red-state (the state of interest in most experiments employing PCFPs).

### The crystal structure of mEos4b-L93M reveals a water cavity near the chromophore

As a first step towards further improvements, we aimed to understand the rationale behind the observed accelerated maturation for mEos4b-L93M. To this aim, we solved the structure of mEos4b-L93M (PDB: 9GVR) by X-ray crystallography, and compared it with the previously published structure of mEos4b (PDB ID: 6GOY).^46^ The only significant difference (beside the L93M mutation) was the presence of an extra water molecule in the vicinity of the chromophore, that is well ordered as evidenced by the clear electron density map at this position (Figure 5A). This water molecule (W1 hereafter) is within hydrogen-bonding distance of the sulphur atom of M93 and the hydroxyl group of T59. It also interacts with the side chain guanidinium group of R91, thereby providing a partner that modulates the interaction of R91 with the chromophore and the hydrogen bond with the side chain of T59 observed in mEos4b (Figure 5B).

**Figure 5.**
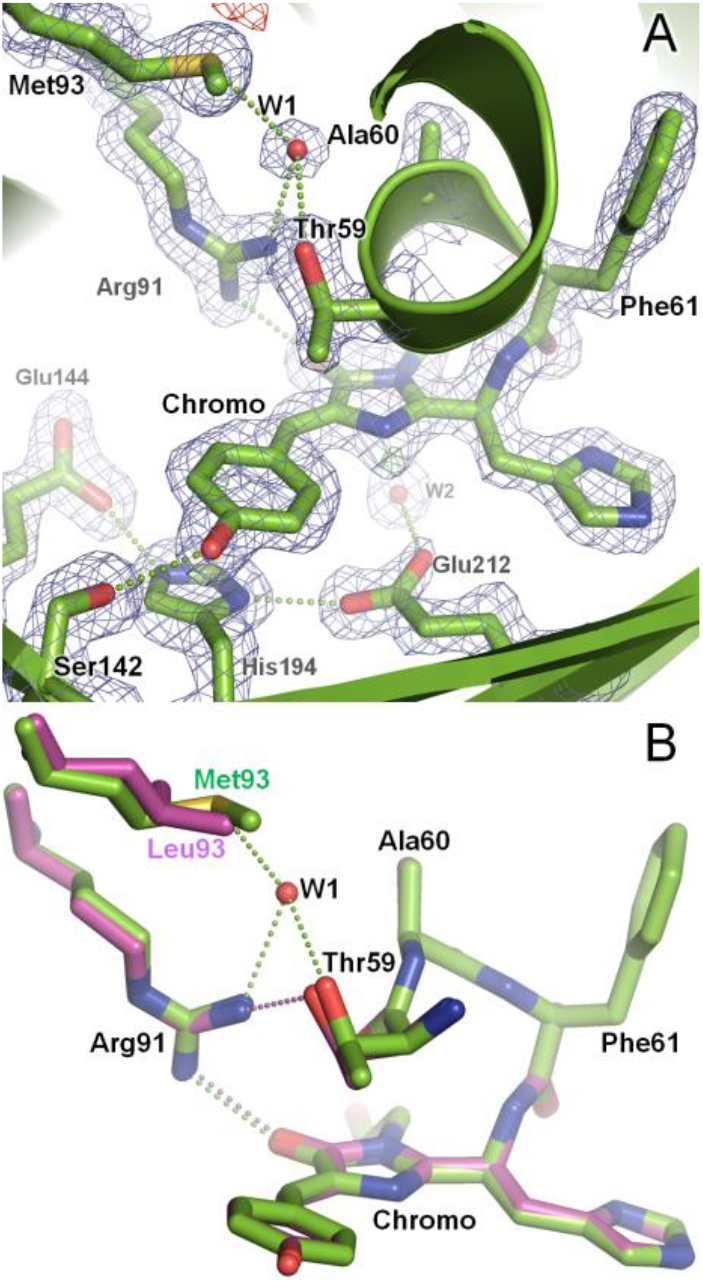
Crystallographic structure of mEos4b-L93M. (A) The chromophore and its environment are shown with green carbons, contoured by 2F_obs_-F_calc_ electron density map (1.5 σ) and F_obs_-F_calc_ difference electron density map (±3.0 σ). The mutation L93M allows the stabilization of a new water molecule, labelled W1. Hydrogen bonds (≤ 3.3 Å) are depicted as green and purple dashed lines for mEos4b-L93M and mEos4b, respectively. (B) Compared to its parent mEos4b, shown with purple carbons, the interaction between W1 and R91 relocalises this arginine relative to the chromophore and breaks its hydrogen bond with the side chain of T59 (from 3 Å to 3.6 Å).

### Fast maturing variants mEos4Fast1 and mEos4Fast2

Aligning the 3D structures of the fast maturing and bright fluorescent proteins mNeonGreen (PDB: 5LTR) and mScarlet3 (PDB: 7ZCT) with that of mEos4b (PDB: 6GOY) (Figure S4), we observed a hydrogen-bonding interaction between the side chains at position 60 and R91 in the 2 fast maturing proteins. This H-bond cannot be formed in mEos4b, which harbours an alanine at position 60. We hypothesized that this interaction might play a role in positioning the key residue R91 so as to facilitate rapid maturation. In addition, an exploration of all fluorescent proteins with reported maturation times in FPbase revealed that the position equivalent to A60 is commonly occupied by glutamine, aspartate, serine or threonine (Figure S5). Based on our structural data, we speculated that this side chain may occupy the space of the water molecule observed for mEos4b-L93M and provide a hydrogen bonding partner to R91. Thus, we generated and tested mEos4b mutations of A60 to either aspartate, glutamine, serine or threonine. In addition we also mutated A60 to histidine and proline, like in the fast-maturing FPs mNeonGreen^34^ and Gamillus,^47^ respectively. Finally, we tested the A60K mutation to explore if a long basic but flexible sidechain could still adapt. Among these mutants, only the A60Q variant yielded a bright fluorescent protein with a significantly (∼9-fold) improved maturation speed compared to the mEos4b parent (Figure 6).

**Figure 6.**
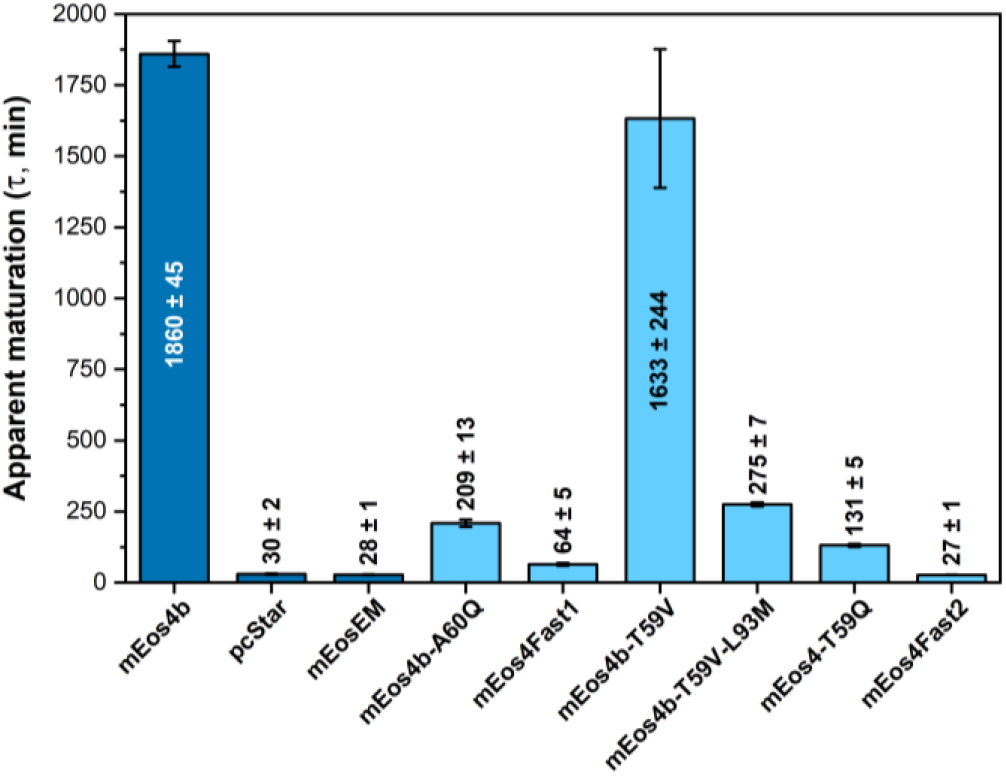
Comparison of time constants of apparent maturation (folding + cyclization + dehydration + oxidation) for mEos4b and selected derivatives (dark blue) as compared to variants that led to the creation of mEos4Fast1 and mEos4Fast2 (light blue). Error bars are standard deviations of time constants extracted from three separate measurements.

Introducing the single A60Q mutation resulted in maturation times of 209 ± 13 min for mEos4b-A60Q and 64 ± 5 min for mEos4b-A60Q-L93M. We named this fast maturing double mutant mEos4Fast1. Despite our efforts, we were unable to crystallize mEos4Fast1, so we used AlphaFold 3^43^ and molecular dynamics (MD) simulations in the Phenix package^44^ to generate structural models of this variant (Figure S6). In both computed structural models, the amide group of Q60 replaces the water molecule W1 observed in the mEos4b-L93M structure, leading to a similar repositioning of the R91 side chain relative to T59.

We also explored a glutamine mutations at position 59, where T59 forms a hydrogen bond with either the water molecule or the guanidinium group of R91 in the mEos4b-L93M and mEos4Fast1 mutants. We prepared the two mutants, mEos4b-T59Q and mEos4b-L93M-T59Q. In addition, we generated mEos4b-T59V and mEos4b-L93M-T59V as negative controls since valine is isosteric with threonine, but lacks a polar side chain. As expected, the apparent maturation of mEos4b-T59V and mEos4b-T59V-L93M was slow (Figure 6).

In contrast, the T59Q mutation significantly improved the apparent maturation time, particularly in the variant mEos4b-T59Q-L93M, which we renamed mEos4Fast2. This protein outperforms any other tested mEos variants and challenges the very fast maturing mEosEM and pcStar (Figure 6). We subsequently proceeded with the biochemical and photophysical characterization of mEos4Fast1 and mEos4Fast2.

### Photophysical and biochemical properties of the new mutants

To investigate whether our fast maturing mEos4Fast1 and mEos4Fast2 variants retained the photophysical and biochemical properties of their mEos4b parent, we compared the *in vitro* fluorescence properties of mEos4b-L93M, mEos4Fast1 and mEos4Fast2 with those of the latest mEos variants, namely mEos4b, pcStar and mEosEM (Table 1).

**Table 1.** Photophysical parameters of the members of the mEos family in their green (G) and red (R) species at pH 7.5 measure at the ensemble level. (^#^value reported in FPbase)

| Protein | $\lambda_{\text{ex, max}}$<br>(nm) | $\lambda_{\text{em, max}}$<br>(nm) | $\epsilon$ ($\text{M}^{-1}\cdot\text{cm}^{-1}$) | QY | Bright. | pKa |
| --- | --- | --- | --- | --- | --- | --- |
| mEos4b (G) | 504 | 518 | 97970 | 0.67 | 65.64 | 5.5<sup>#</sup> |
| mEos4b (R) | 569 | 580 | 56130 | 0.52 | 29.19 | 5.8<sup>#</sup> |
| pcStar (G) | 504 | 516 | 80910 | 0.61 | 49.36 | 5.5<sup>#</sup> |
| pcStar (R) | 569 | 580 | 47940 | 0.29 | 13.90 | 6.1<sup>#</sup> |
| mEosEM (G) | 504 | 515 | 88280 | 0.69 | 60.91 | 5.6<sup>#</sup> |
| mEosEM (R) | 570 | 581 | 51090 | 0.33 | 16.86 | 6.5<sup>#</sup> |
| mEos4b-L93M (G) | 504 | 515 | 77120 | 0.68 | 50.44 | 5.6 |
| mEos4b-L93M (R) | 569 | 580 | 57200 | 0.43 | 24.60 | 6.3 |
| mEos4Fast1 (G) | 505 | 517 | 93170 | 0.78 | 71.37 | 5.6 |
| mEos4Fast1 (R) | 570 | 580 | 58450 | 0.50 | 29.22 | 6.7 |
| mEos4Fast2 (G) | 499 | 513 | 78020 | 0.82 | 63.98 | 4.9 |
| mEos4Fast2 (R) | 565 | 580 | 43290 | 0.37 | 16.02 | 5.4 |

At the ensemble level, both mEos4Fast1 and mEos4Fast2 exhibit brightness in their green fluorescent form comparable to mEos4b and mEosEM, and significantly higher than pcStar. In its red form, mEos4Fast1 is as bright as mEos4b and significantly brighter than pcStar and mEosEM, while for red mEos4Fast2 similar values are found. Of note, mEos4Fast2 features a surprisingly low pKa both in its green and red forms. A low pKa is advantageous for ensemble brightness in a cellular context, notably in acidic compartments, but can also be problematic for green-to-red photoconversion as it requires more phototoxic 405-nm light to excite the neutral chromophore. While green chromophores of all studied PCFPs are nearly entirely in their anionic (fluorescent) form at physiological pH (7.2), the red chromophores of pcStar and mEosEM are 87% and 76% in the anionic form, respectively, compared to 97.5% for mEos4Fast2.

At the single-molecule level, mEos4b, mEos4Fast1 and mEos4Fast2 were compared for their blinking propensities, on-times, off-times, photon budget and photostability *in vitro* using polyacrylamide gel embedding (Figure S7). The proteins were essentially indistinguishable from each other in terms of these properties, although we noticed that red mEos4Fast2 was slightly less bright at the single molecule level and exhibited slightly shorter on-times and longer off-times than the other variants. This possibly suggests facilitated *cis-trans* isomerization of the mEos4Fast2 chromophore as a source of blinking. Overall, we concluded that the faster maturation of mEos4Fast1 and mEos4Fast2 did not degrade the excellent single-molecule photophysical properties of mEos4b.

Finally, we ensured that our fast-maturing mEos4b variants retained their properties in common fixation reagents used in cell imaging, such as paraformaldehyde (PFA) and glutaraldehyde (GA).^48^ While mEos4Fast1 showed moderate stability in the presence of these reagents, mEos4Fast2 emerged as the best fluorescent protein we tested, retaining 70-80% of its fluorescence after fixation (Figure S8).

### Effect of faster maturation on apparent cellular brightness

The apparent cellular brightness and maturation speed of mEos4Fast1 and mEos4Fast2 were compared to those of their parent mEos4b in prokaryotic and eukaryotic cells. For prokaryote expression, cells were grown, plated and imaged regularly in fluorescence mode. After IPTG-induction, a striking difference in green fluorescence signal was observed between the two fast variants and mEos4b, with mEos4Fast2 becoming quickly brighter than mEos4Fast1, which in turn was significantly brighter than mEos4b (Figure S9). For eukaryote expression, U2OS cells were transfected with the bicistronic constructs mEos4b-2a-mCherry, mEos4Fast1-2a-mCherry or mEos4Fast2-2a-mCherry. mCherry, connected to the target protein via the small viral T2A self-cleavable linker, served to standardize the different transfection/expression levels in different cells. 3 h after transfection, cells were imaged every 30 min. The resulting time-lapse (Figure 7A, Figure S10 and supplementary video) clearly shows that the green signal from mEos4Fast2 (and to a lesser extent mEos4Fast1) appears much earlier than that of mEos4b. 40 hours after transfection, mEos4Fast2 (and to a lesser extent, mEos4Fast1) continued to show a much brighter signal than mEos4b (Figure 7B), similarly to what we observed in living *E. coli* cells. This observation was confirmed by flow cytometry analysis (Figure 7C, Figure S11).

**Figure 7.**
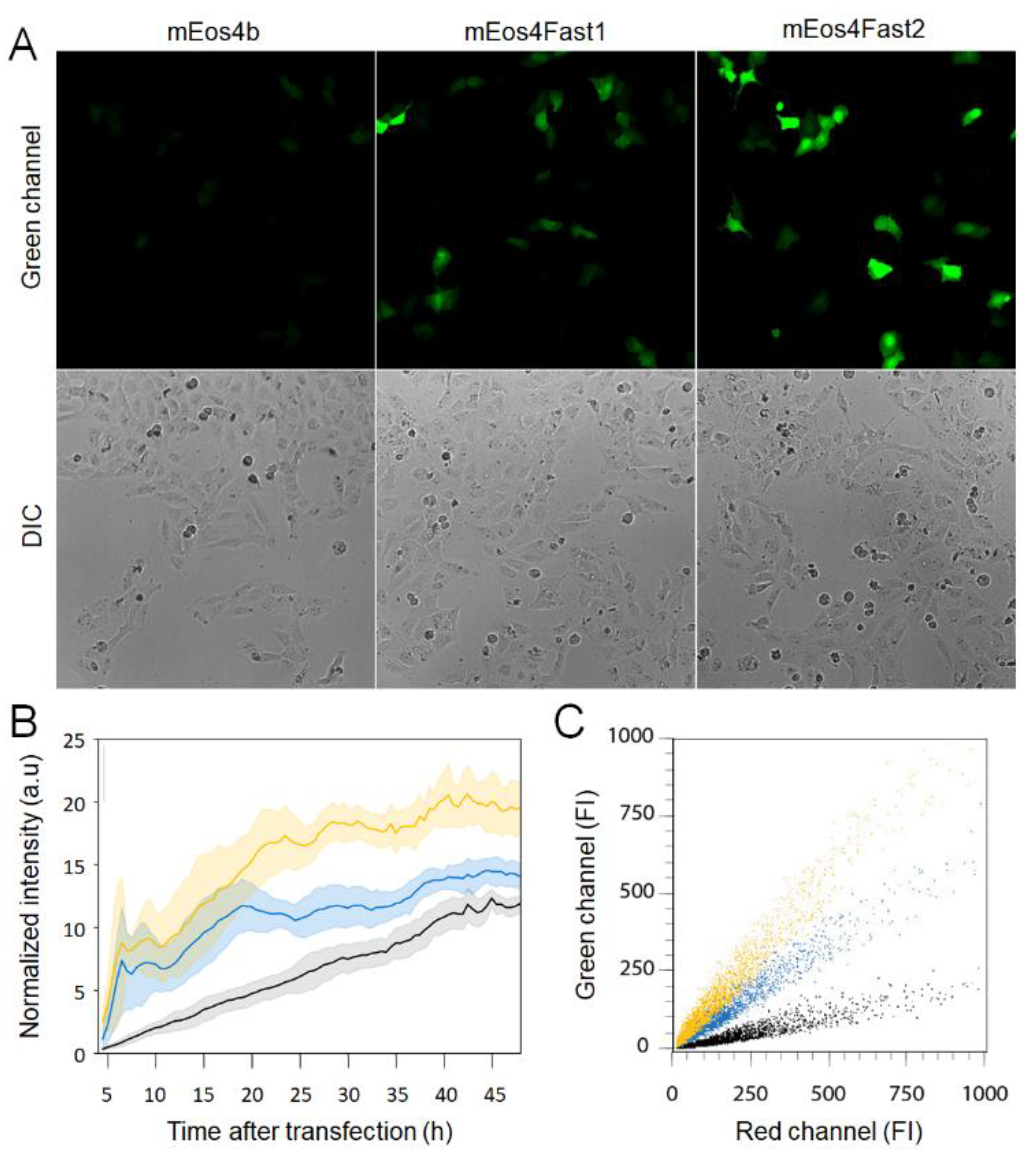
Maturation of mEos4b, mEos4Fast1 and mEos4Fast2 in bicistronic constructs with mCherry in U2OS cells. (A) Representative microscopic images of live U2OS cells transfected with each construct and observed 16 h after transfection. The green channel is normalised by the red channel of mCherry and each field of view contains a comparable number of cells as testified by images recorded in the DIC channel. (B) Mean fluorescence signal evolution for mEos4b (black), mEos4Fast1 (blue) and mEos4Fast2 (yellow) from live U2OS cells in five different field of views (n=5). The green channel is normalised by the signal of mCherry at the end of the kinetics. (C) Flow cytometry analysis of cells 16 h after transfection (10 000 cells per construct). Dot plot of mEos4 (green) versus mCherry (red) fluorescence is shownfor each cell population (10 000 cells per construct).FI: Fluorescence Intensity.

We further questioned why mEos4Fast2 exhibits a higher fluorescence in cells and better resistance to PFA fixation, despite not having a higher molecular brightness measured on purified proteins (Table 1). Part of the explanation lies in the 5-nm blue shift in its absorption spectrum as compared to mEos4b, with an absorption maximum at 499 nm better matching excitation by typical 488 nm light. This hypsochromic shift is also observed in the red form (Table 1, Figure S12), well suited for excitation at 561 nm. Other explanations could involve better expression, solubility or stability of mEos4Fast2 in cells, reduced cytotoxic effects, or better maturation yield.

To verify these hypotheses, we performed a series of control experiments. We initially quantified protein expression levels and maturation by extracting proteins from IPTG-induced *E. coli* cultures (12 hours post-induction), followed by SDS-PAGE analysis and UV-visible spectrophotometry (Figure S13). The results revealed that mEos4Fast2 exhibited the highest maturation efficiency, despite similar protein expression levels for mEos4b and mEos4Fast2, and lower expression for mEos4Fast1. This more efficient maturation may explain the increased brightness observed for mEos4Fast2 both in prokaryotic and eukaryotic cells. Subsequently, we assessed potential cytotoxic effects by measuring the 600-nm turbidity of bacterial cultures before and 12 hours after IPTG induction (Figure S14), but no significant differences in growth ratios among the three proteins could be noticed. Lastly, we evaluated thermal stability by incubating purified mEos4b, pcStar, mEosEM, mEos4Fast1, and mEos4Fast2 at temperatures ranging from 66°C to 80°C and measuring residual fluorescence (Figure S15). mEos4Fast2 ranked second in thermal resistance, very close to mEosEM, suggesting that its structural robustness contributes to the stability of mEos4Fast2 in cells. These combined factors, (efficient and faster maturation, protein stability and better resistance to fixation reagents) account for the consistently higher brightness of mEos4Fast2 observed in living and fixed cells.

### Slow cyclization causes slow maturation of mEos4b

We further investigated whether pre-oxidation or post-oxidation events are responsible for the extremely slow maturation of mEos4b. Our aerobic maturation assay essentially measures the combination of all the processes involved in folding and maturation. We modified the assay to work under anaerobic conditions (see methods and Figure S16), to ensure that the protein is produced and subsequently folds but, in the absence of oxygen, maturation of its chromophore stalls at the pre-oxidation step(s). The fluorescence increase obtained upon subsequent exposure to oxygen then reflects the kinetics of the oxidation and (eventual) post-oxidation events. Strikingly, under these conditions, the rise in fluorescence was fast, on the order of 10 minute, for all tested mEos variants. This is in contrast with the results obtained from the aerobic maturation assay (Figure 8). These results thus strongly suggest that for mEos4b and other slow maturing mEos derivatives pre-oxidation events are rate limiting, being considerably slower than the rest of the maturation process.

**Figure 8.**
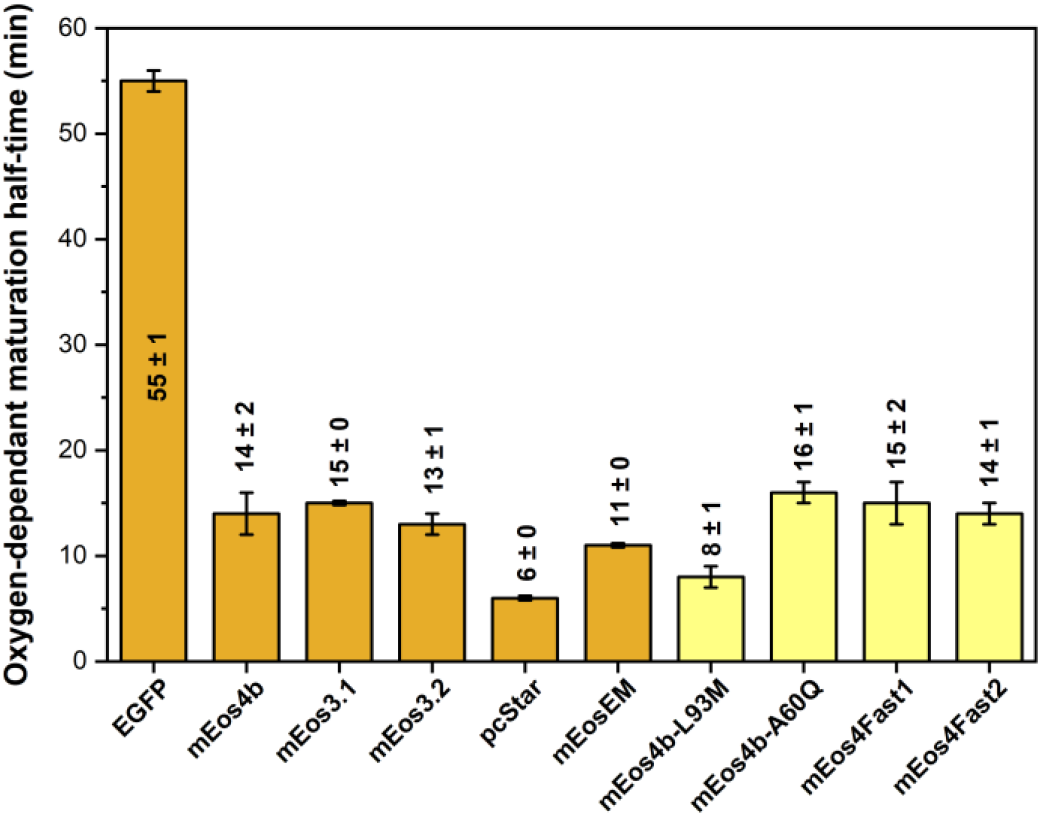
Comparison of half-times for oxygen-dependent chromophore forming step(s). Results are presented for EGFP (control), selected members of the mEos4b family (orange), and mEos4Fast1 and mEos4Fast2, as well as some initial variants generated for this study (yellow). Error bars are standard deviations of time constants extracted from three separate measurements.

Next, we aimed to identify or narrow down which steps of the pre-oxidation events is actually rate-limiting during mEos4b maturation: protein folding, cyclization, or dehydration for pathway A (Figure 1). First, protein folding can be excluded from this list because our NMR data showed that the slowly maturing species, detected for fresh mEos4b samples, displays only minor chemical shift differences compared to mature mEos4b (Figure 2). In the case of an unfolded, or only partially folded protein, much larger chemical shift differences would be expected. Among the two remaining possibilities, we speculate that cyclisation is the rate-limiting step. This hypothesis is supported by the fact that R91 has been shown to be a key residue to catalyse cyclisation, while our findings indicate that mutations modulating the interaction of R91 with the chromophore are essential to accelerate maturation. However, it cannot be excluded that dehydration could also be rate limiting if pathway A is followed, as the enolate form of the chromophore, promoted by R91, might be required for this step.

## Discussion

Our study tackles the significant challenge posed by the slow maturation of mEos4b, a widely used PCFP. This slow maturation compromises effective labelling efficiency and may hinder real-time tracking of dynamic processes. We developed a strategy to generate fast-maturing variants of mEos4b, which are functional both in bacteria and mammalian cells. Importantly, this accelerated maturation does not compromise the biochemical and photophysical properties of the protein. The variants retain the favourable properties of the parent mEos4b at both the ensemble and single-molecule levels, ensuring their reliability and effectiveness in advanced fluorescence microscopy.

Our work provides a comprehensive comparison of the maturation rates among various mEos-derived PCFPs, revealing significant differences between them. Although absolute values may be influenced by the methodology used, our measurements offer a reliable comparison. When maturation speed is critical, we recommend that users select mEosEM, pcStar or one of our fast-maturing mEos4b variants for their experiments. All four PCFPs exhibit relatively similar properties at the ensemble and single molecule levels, with mEosEM and mEos4Fast2 having particular advantages. Utilizing these fast-maturing PCFPs should enable real-time visualization of dynamic events at higher resolution, which is essential for studying proteins with short half-lives, or observing protein synthesis, modification, and degradation without the delays associated with slower-maturing chromophores.

Significant acceleration of mEos4b maturation was achieved by modifying the immediate environment of R91, a residue known to play a key role in catalysing maturation. In the absence of a structure for the pre-cyclized chromophore, the exact mechanism by which the introduced mutations alter the interaction of R91 with the carbonyl group of G64 (the third amino acid of the chromophore) remains to be clarified. We hypothesize that a slight repositioning of R91 and/or changes in the charge distribution at the guanidinium group could favour the cyclisation step, and/or possibly the dehydration step during maturation. High-level theoretical calculations may provide additional insights into the exact mechanisms involved.

Our finding that the cyclization step may be rate-limiting during mEos4b maturation is in stark contrast with previous observations in fast-maturing FPs,^21, 28, 49^ where cyclization occurs rapidly, and the rate-limiting steps are oxygen-dependent processes. This difference is confirmed by our EGFP maturation data measured under anaerobic conditions (Figure 4), revealing a much slower oxidation step compared to all mEos-derived variants. This suggests that the free energy barriers associated with the various maturation steps strongly differ between hydrozoan and anthozoan proteins. Given the relatively low sequence identity (∼30%) between FPs from the two families, this difference may not be so surprising. However, further investigations will be necessary to explore this issue in greater detail.

In PALM, sptPALM or MINFLUX, fast-maturing mEos variants may be indispensable for some experiments, both in fixed or live cells. First, a swift fluorescence onset is essential to rapidly capture at the nanoscale molecular patterns triggered by e.g. drug administration. Second, fast and efficient maturation can be particularly crucial, depending on target-protein turnover, to maximise effective labelling efficiency in counting experiments aimed at e.g. measuring protein complex stoichiometries. Third, achieving sufficient labelling density is central to the reconstruction of truly high-resolution nanoscopy images and for providing a faithful representation of the spatial distribution of low-abundance proteins.

As compared to mEos4Fast1, mEos4Fast2 matures even faster and has a lower pKa, leading to higher overall brightness in live cells. Furthermore, mEos4Fast2 displays strong resistance to chemical fixation. Thus this novel mEos variant is expected to perform exceptionally well in live-or fixed-cell applications, making it a marker of choice for pulse-chase ensemble or single-molecule based super-resolution imaging, including correlative studies. mEos4Fast1, on the other hand appears better suited for single-particle tracking experiments, for which a reduced blinking propensity is crucial for accurate tracking.

## Methods

### Cloning

All the proteins studied were cloned in a modified pET28a vector carrying the gene for kanamycin resistance with the exception of mEos2, mEos3.1 and mEos3.2 that we already had cloned in pRSET-A vector carrying the gene for ampicillin resistance.

### Mutagenesis

Single point mutants were prepared in house using the Q5^®^ Site-directed mutagenesis kit (New England Biolabs) and primers whose list can be found in Table S2.

### Mammalian bicistronic constructs

For enhanced mammalian expression, we optimized the mEos4b sequence by incorporating a Kozak sequence and N-terminal (N-ter) and C-terminal (C-ter) sequences from EGFP, a widely adopted strategy^50^. Through PCR amplification, we substituted the two N-terminal amino acids of mEos4b (MS) with the 11-residue GFP-type N-terminus sequence (MVSKGEEDNMA). Concurrently, we ensured placement of the Kozak sequence (GCCACCATGG) between the *Nhe*I site and overlapping the initiation codon of the mEos4b coding sequence. Similarly, the C-terminal sequence of mEos4b (DNARR) was replaced with the 7-residue GFP-type C-terminus sequence (GMDELYK) in a symmetrical manner. A 24-residue linker (GSGEGRGSLLTCGDVEENPGPRSL) was also inserted immediately after mEos4b sequence and overlapping the *Age*I restriction site. This linker codes for the underlined 18-residue 2A peptide from the *Thosea asigna* virus capsid protein, self-cleaving between the last two amino acids (proline and glycine) and ensuring an equimolar translation of the two fluorescent proteins. This sequence was cloned between the *Nhe*I and *Age*I restriction sites in the MCS, upstream mCherry in the mammalian plasmid pmCherry_N1 (Clontech) and mutations T59Q, A60Q and L93M were then added to mEos4b, resulting in the three bicistronic constructs mEos4b-2a-mCherry_N1, mEos4Fast1-2a-mCherry_N1 and mEos4Fast2-2a-mCherry_N1.

### Protein production and purification

*Escherichia coli* BL21(DE3) cells were transformed with a plasmid carrying the target protein. Cells were adapted from rich LB medium to M9 minimal medium isotopically enriched with ^15^N NH_4_Cl (1 g/L) and ^13^C-glucose (2 g/L) in two steps over 24 h. For proteins prepared for X-ray crystallography, the adaptation from LB to M9 minimal media was not necessary; therefore, all steps of cells growth were performed in LB medium. Cells were grown at 37°C until the turbidity measured at 600 nm reached >0.5. Protein over-expression was induced by adding 1 mM IPTG (isopropyl β-D-thiogalactopyranoside). After overnight induction at 20 °C, cells were harvested and sonicated in 20 mM HEPES buffer at pH 7.5 supplemented with 150 mM NaCl and a cOmplete™ EDTA-Free (Roche) tablet. The lysate was centrifuged at 46000g for 40 min. The supernatant was collected and passed through a Ni-NTA (Qiagen) column pre-equilibrated with the buffer mentioned before. Protein elution was performed with 500 mM imidazole. Size exclusion chromatography was performed with a S75 (Cytiva) column equilibrated with 20 mM HEPES pH 7.5, 150 mM NaCl Buffer. Elution fractions containing the protein were dialyzed against the 20 mM HEPES buffer at pH 7.5. Purified samples were stored at -80°C until needed.

### NMR experiments

NMR samples were prepared in 5 mm Shigemi NMR tubes, containing 100-200 μM of protein in 300 μL of buffer solution with 5 % (v/v) D_2_O, filled. NMR experiments were performed on a Bruker Avance III-HD spectrometer (850 MHz) equipped with a triple-resonance cryo-probe and pulsed z-field gradients. Unless otherwise specified, all measurements were performed at 35°C. Data were processed using TopSpin v3.5 (Bruker BioSpin). Data analysis was performed with the CCPNMR v2 software. 2D ^1^H-^15^N amide backbone correlation spectra were recorded using a BEST-TROSY pulse sequence^51^ with the ^15^N carrier centred at 120 ppm, and the ^1^H band selective pulses covering a band width of 8.7 ± 2.5 ppm.

### Fluorescence-based maturation assays

Maturation assay performed in aerobic condition consisted of similar protocol for protein production described above, consisting of transformation, cell growth in LB medium and induction by IPTG at 37°C for 1-2 hrs. A cocktail of antibiotics containing chloramphenicol (final concentration 0.17 mg/mL) and tetracycline (final concentration 0.05 mg/mL) were then added to the media containing the cells to stop fresh protein production. 100 μL of the solution was transferred in well(s) of 96-well plate. Fluorescence signal was measured in a BioTek microplate reader. Fluorescence signal was excited at 470 nm, and the emission data were integrated between 495-535 nm. The signal corresponding to LB medium only were subtracted from each dataset. The data were then baseline corrected for any turbidity variations measured at 600 nm during the measurement process. This ensured no data artefacts introduced due to delay in antibiotic functioning resulting in residual cell division and cell death. Homemade python scripts were used to perform data analyses. The data were plotted as a function of time to follow the maturation process. Most datasets fit (exceptions are mentioned in results section) to a mono-exponential equation. The maturation time reported in this study is the mono-exponential time-constant obtained from the fitting. All experiments were triplicated. In order to perform the maturation assay in anaerobic conditions the following adjustments to the assay were brought into. The cells were grown overnight in presence of oxygen in LB medium at 37°C. Following which a subculture was done by diluting the cells 100-times in Terrific Broth medium supplemented with 50 mM MOPS/KOH at pH 7.1, 0.5 % D-glucose and necessary antibiotic (Kanamycin/Ampicillin). Once the bacterial turbidity reached 0.5-0.6, cultures were transferred into glass containers and sealed with caps containing rubber septum. After a 20 min incubation period at 37°C (to ensure all the residual oxygen is consumed by the cells), the culture was supplemented with ammonium ferric citrate (40 μM), L-cysteine (400 μM), sodium fumarate (25 mM), vitamin cocktail mix and IPTG by mode of injection through the rubber septum. Following a 2-h induction at 37°C, the antibiotic cocktail containing chloramphenicol (final concentration 0.17 mg/mL) and tetracycline (final concentration 0.05 mg/mL) were added to the culture. The culture bottles were transferred to 20°C for overnight. Following morning, the bottles were opened and 100 μL of culture from each bottle were transferred to 96-well plates for fluorescence measurements. The time differences between opening each bottle and starting of the measurement (typically 2 min) were taken into account during data analyses. A graphical illustration of the assays is shown in Figure S12.

### Ensemble spectroscopic properties and pKa

#### Extinction co-efficient

The molar extinction co-efficient of the green-form of all PCFPs reported in the table 1 was determined using the ward method.^52^ The extinction co-efficient of the free GFP-like chromophore is known to be 44000 M^-1^.cm^-1^ at 447 nm. Comparing the absorption spectra of the native proteins at pH 7.5 and fully denatured proteins with still intact chromophore after gradual addition of 1 M NaOH, the molar extinction co-efficient at their absorption maxima were calculated. The molar extinction co-efficient of the red form of the PCFPs were obtained by comparing the decrease in absorption in the green form and increase in absorption of the red form.^53^ PCFPs were photoconverted in a cuvette (with a starting green-form OD 0.5-0.7) with 405 nm laser of power density 250 mW/cm^2^.

#### Quantum yield

The quantum yield values were measured by comparing with known standards, Fluorescein in 0.1 M NaOH solution for the green forms and Rhodamine 6G in ethanol solution for the red forms. The fluorescence spectra of the green- and red-forms were obtained using an excitation laser of 473 and 532 nm respectively.

#### pKa

Solution of the green and red forms of the PCFPs were prepared at different pH values ranging from 4.2 to 9. From the absorption spectra of the protein solutions the amplitude of the absorption peaks corresponding to either the neutral or the anionic form were plotted as a function of pH. pKa value were obtained by fitting the following equation.

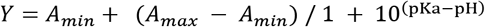

### Crystallization and crystallographic characterization

Purified mEos4b-L93M was concentrated by ultrafiltration and equilibrated in buffer solution (50 mM HEPES pH 7.5). Crystals were obtained by the hanging-drop vapour diffusion method at 20 °C. Briefly, the protein (15 mg/ml) and precipitant solution (0.1 M HEPES pH 8.5, 32 % PEG 1000) were mixed 1:1, yielding 2-μl drops that were placed over a 1 ml well containing the precipitant solution. Needle-shaped crystals appeared within 7 days. Prior to data collection, crystals of mEos4b-L93M were cryoprotected by a short soak in the mother liquor supplemented with 20 % glycerol, followed by flash-cooling in liquid nitrogen. X-ray diffraction data sets were collected at 100K at the European Synchrotron Radiation Facility (ESRF) on the beamline ID30-A3/MASSIF-3,^54^ equipped with an Eiger1 × 4M detector and a wavelength set to 0.9677 Å. Data were processed via the autoPROC pipeline^55^ using XDS^56^ and POINTLESS^57^ to integrate and merge data in the P2_1_2_1_2_1_ spacegroup and AIMLESS^58^ for scaling. The structure was phased by the molecular replacement method using as a starting model the X-ray structure of mEos4b (PDB ID: 6GOY) and the program Phaser.MRage.^59^ Model building was performed with *Coot*.^60^ Refinement and map calculations were performed using Phenix^44^ at 1.86 Å resolution. Data collection and refinement statistics are provided in Table S3. Figures were produced using PyMOL.^61^

### Apparent brightness in bacteria

*E. coli* cells were transformed with plasmids containing the coding sequence of either mEos4b, mEos4Fast1 or mEos4Fast2 under the control of the T7 promoter. Cultures were let to grow at 37°C until reaching a turbidity of 0.5 at 600 nm. Cultures were then induced with 10 mM IPTG, immediately plated on a LB-agar gelose and let at room temperature in the dark. From 4 hours after plating, photographs were taken regularly under a blue light transilluminator, during 24 hours.

### Transfection

U2OS cells were seeded at a density of 0.4 × 10^6^ cells per well in six-well plates containing growth medium (phenol red-free DMEM supplemented with 10% FBS). The following day, the cells were transfected with bicistronic constructs, including mEos4b-2A-mCherry, mEos4Fast1-2A-mCherry, or mEos4Fast2-2A-mCherry, using the Fugene HD reagent (Promega) as per the manufacturer’s instructions. The transfected cells were then incubated at 37°C with 5% CO2 for up to 40 hours, depending on the subsequent analysis method.

### Apparent brightness in human cells

Fluorescence microscopy experiments were conducted using U2OS cells transfected with either the mEos4b-2a-mCherry, the mEos4Fast1-2a-mCherry or the mEos4Fast2-2a-mCherry construct. Following transfection, cells were cultured in FluoroBrite™ DMEM (Gibco) medium to minimize autofluorescence, supplemented with 10% FBS, glutamax (Gibco) and MEM NEAA (Gibco). Cells were then observed under a spinning disk microscope, Olympus IX81, equipped with a Yokogawa CSU-X1 confocal head, a motorized stage and an incubation chamber (Okolab), which provides 5% CO_2_ atmosphere at 37°C to maintain a stable pH throughout the live cell imaging. Time-lapse acquisition commenced as early as 4 h post-transfection, with the time interval between frames set to 30 minutes. The sample was excited with a 488-nm laser at 0.3 mW (green channel, Eos variants) and a 561-nm laser at 0.13 mW (red channel, mCherry). Laser powers were measured at the sample level (Ilas2 laser bench, GATACA systems). Fluorescence signal was collected with a 10X 0.25 NA air-objective and filtered with individual band-pass filters Semrock FF02-520/28 and Chroma ET600/50m, for green and red channels respectively, before being detected by an EMCCD camera (Andor iXon Ultra). Fluorescence kinetics were monitored within cells for a period of 44 h. The green fluorescence signal was normalized to the red signal originating from mCherry, allowing for accurate quantification and comparison of fluorescence levels between experimental conditions.

### Fixation resistance assay

Fluorescence conservation ability of the PCFPs in presence of fixating agents PFA and GA were tested based on methods demonstrated before.^48^ In brief, purified proteins were diluted in PBS (pH 7.2), and incubated for 30 min at 37°C either without any fixative agents or with A) 4% PFA and B) 4% PFA + 0.2% GA. Following which fluorescence intensity were measured in a BioTek microplate reader. Measurements were also performed on transfected U2OS cells. Twenty-four hours after transfection, the fluorescence of the same harvested cells sample was analysed by flow cytometry, before fixation (live cells), after 30 minutes of treatment with 4% PFA or with 4% PFA + 0.2% GA. Fixation resistance was calculated based on the ratio of fluorescence intensity of samples incubated with fixatives to the corresponding controls (without fixatives). The data is presented in Figure S8.

### Flow cytometry

Following transfection, the cells were maintained in culture for 8, 16, 24 or 40 hours at 37°C with 5% CO_2_. After the designated incubation period, the cells were harvested using 0.25% trypsin/EDTA. The trypsinisation process was halted by adding medium containing 10% FBS. The cells were then pelleted by centrifugation at 300 g for 5 minutes, washed, and resuspended in PBS for flow cytometry analysis. Flow cytometry was performed using a MACSQuant VYB Cytometer (Miltenyi Biotec), with a minimum of 10,000 events collected within the analysis gate. mCherry and green state mEos4 fluorescence were acquired simultaneously. Data were analysed using MACSQuantify software.

### Simulation of the A60Q mutation

We employed AlphaFold 3,^43^ a cutting-edge protein structure prediction tool (available at https://golgi.sandbox.google.com), to simulate the 3D structure of mEos4b carrying the A60Q and L93M mutations, starting from the amino acid sequence. Concurrently, we utilized the dynamics and minimization (phenix.dynamics) module of Phenix version 1.20.1-4487,^44^ with the crystallographic structure of mEos4b-L93M as a starting model in which the A60Q mutation was first introduced and water molecule W1 removed, using Coot v. 0.9.8.93.^60^

### Chromophore and protein yields for mEos4b, mEos4Fast1 and mEos4Fast2

*E. coli* beta-10 (NEB) colonies expressing either mEos4b, mEos4Fast1 or mEos4Fast2 were inoculated in LB medium supplemented with kanamycin. Cultures were grown at 37°C until reaching a turbidity of 0.5 at 600 nm. Protein expression was induced in 4 mL culture with 10 mM IPTG, followed by overnight incubation at 37°C. Cells were harvested, and proteins were extracted using BugBuster Protein Extraction Reagent (MilliporeSigma, Cat. No. 70584-3) according to the manufacturer’s protocol. The extracts were analysed by SDS-PAGE and UV-visible spectrophotometry to determine protein concentration and maturation.

### Single molecule fluorescence measurements

Purified proteins (mEos4b, mEos4Fast1, and mEos4Fast2) were immobilized in a polyacrylamide gel (pH 8) and single molecule imaging was performed using laser excitations at 561 nm (500 W/cm^2^, 70 ms exposure time) and 405 nm (1 W/cm^2^, 8.2 ms exposure time). Fluorescence was collected by a 100X 1.49 NA oil-immersion objective (Olympus) and detected by an EMCCD camera (Evolve 512, Photometrics) and a pixel size of 128 nm/pixel.

Molecules were localized using the ThunderSTORM ImageJ plugin^62^ and localizations were clustered using an in-house MATLAB routine to reconstruct fluorescent time traces belonging to single molecules.^63^ From these fluorescent time traces, off-times, on-times, bleaching-times, number of blinks and photon budget were extracted. All proteins behave similarly under these conditions.

## Conclusions

The development of fast-maturing PCFPs is needed to enhance the quality of data obtained from advanced fluorescence microscopy techniques. In addition to mEosEM and pcStar, our new fast maturating mutants mEos4Fast1 and mEos4Fast2 are expected to be particularly beneficial for PALM and MINFLUX super-resolution microscopy, addressing the limitations of the very slow-maturing mEos2, mEos3.1, mEos3.2 and mEos4b variants and providing a reliable tool for capturing transient and dynamic cellular events

Our study is the first to propose a rational design approach for engineering fast maturing variants of a photoconvertible fluorescent protein. It provides new mechanistic insights into the maturation mechanism of anthozoan FPs. However, further research is needed to fully understand the precise role of the key mutations introduced around R91 in accelerating maturation. Additionally, evaluating the performance of our fast-maturing mEos4b variants across a diverse range of experimental conditions and biological systems will be essential for expanding their applicability. Lastly, applying our strategy to other slow-maturing FPs, including other PCFPs, could advance various fields of biological and biomedical research, enabling faster and more precise cell imaging.

## Author contributions

D.B. and B.B. did the early conceptualization of the project and acquired financial support for it. B.B. performed NMR experiments. V.A. conceived and supervised the project. V.A. and A.M. designed the experiments. I.A. and A.M. produced and purified fluorescent proteins. V.A. designed and performed molecular biology experiments, created mutants and made bacterial expression and stability assays with help from P.T. A.M. performed spectroscopic characterization, ensemble photophysics characterization, maturation assays and kinetics. J.W. made single molecule measurements, J.W. and B.D. made the single molecule data analysis. V.A. crystallized mEos4b-L93M, solved its structure and performed structure dynamics simulations. P.T and P.F. performed cell biology and flow cytometry experiments. O.G. performed mammalian cell imaging with help of P.T. and P.F. and analysed data.

V.A. and M.A. wrote the manuscript with input from all authors. All authors have given approval to the final version of the manuscript.

## Data availability

The data for this manuscript is provided in the main text and ESI.

## Acknowledgements

Financial support from the Agence Nationale de la Recherche (grants no. ANR-20-CE11-0013-01 and ANR-22-CE11-0011-01) (FR2054) is gratefully acknowledged. This work used the platforms of the Grenoble Instruct-ERIC centre (ISBG; UAR 3518 CNRS-CEA-UGA-EMBL) within the Grenoble Partnership for Structural Biology (PSB), supported by FRISBI (ANR-10-INBS-0005-02) and GRAL, financed within the University Grenoble Alpes graduate school (Ecoles Universitaires de Recherche) CBH-EUR-GS (ANR-17-EURE-0003). IBS acknowledges integration into the Interdisciplinary Research Institute of Grenoble (IRIG, CEA).The authors acknowledge Joël Beaudouin for designing the initial contruct of the bicistronic plasmid, Elke De Zitter for collecting X-ray diffraction data, Martin Weik and Ninon Zala for facilitating the access to use the plate reader and Yvain Nicolet for helpful discussions to design the anaerobic maturation assay.

