## Supplementary information for "Decoding mEos4b Day-Long Maturation and Engineering Fast Maturing Variants"

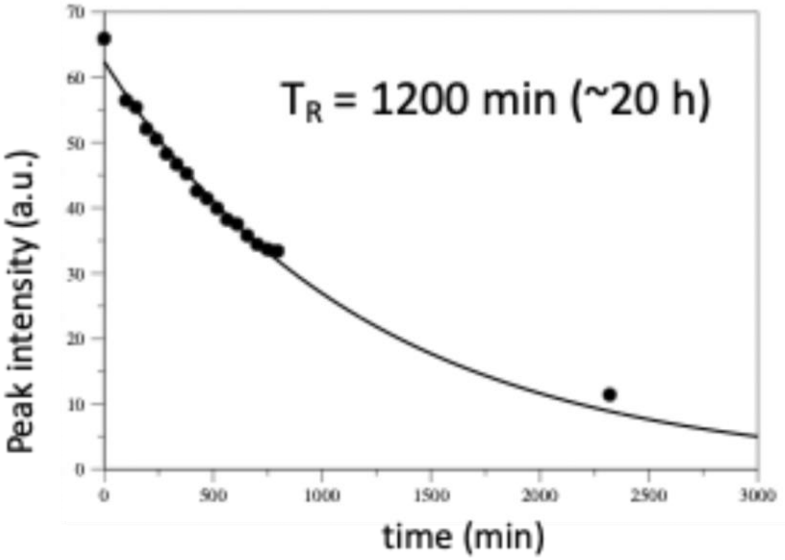


*Figure S1. Kinetics of disappearance of peaks assigned to protein molecules with immature chromophore as observed in $^{1}H$-$^{15}N$ correlation experiments in NMR.*

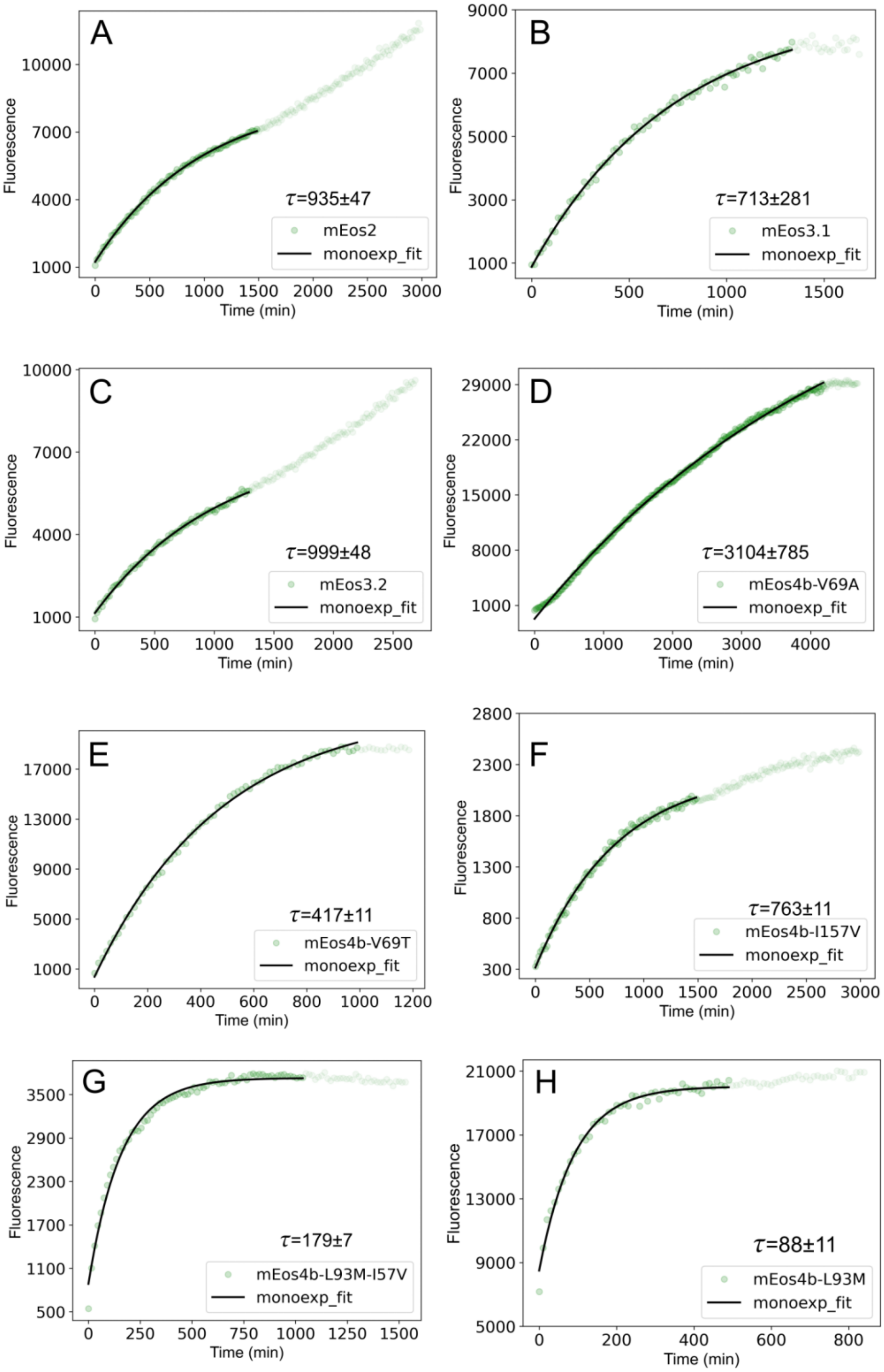


*Figure S2. Fluorescence development traces for the maturation assay of mEos2 (A), mEos3.1 (B), mEos3.2 (C), mEos4b-V69A (D), mEos4b-V69T (E), mEos4b-I157V (F), mEos4b-L93M-I157V (G) and mEos4b-L93M respectively.*

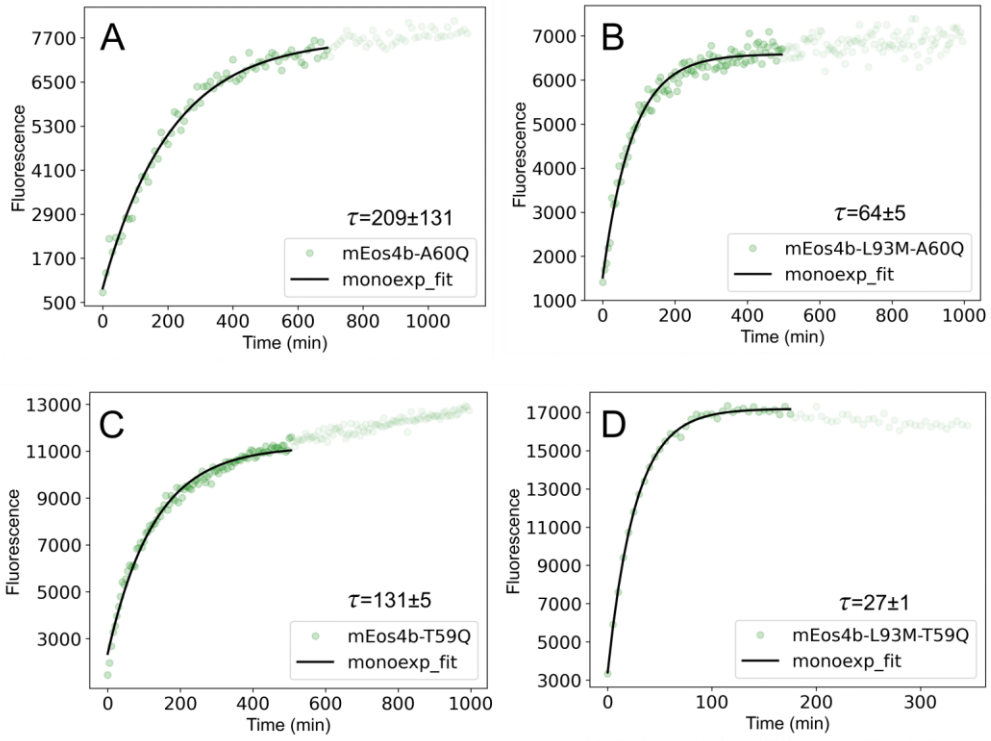


*Figure S3. Fluorescence development traces for the maturation assay of A60Q and T59Q mutants mEos4b-A60Q (A), mEos4b-A60Q-L93M (mEos4Fast1) (B), mEos4b-T59Q (C) and mEos4b-T59Q-L93M (mEos4Fast2) (D).*

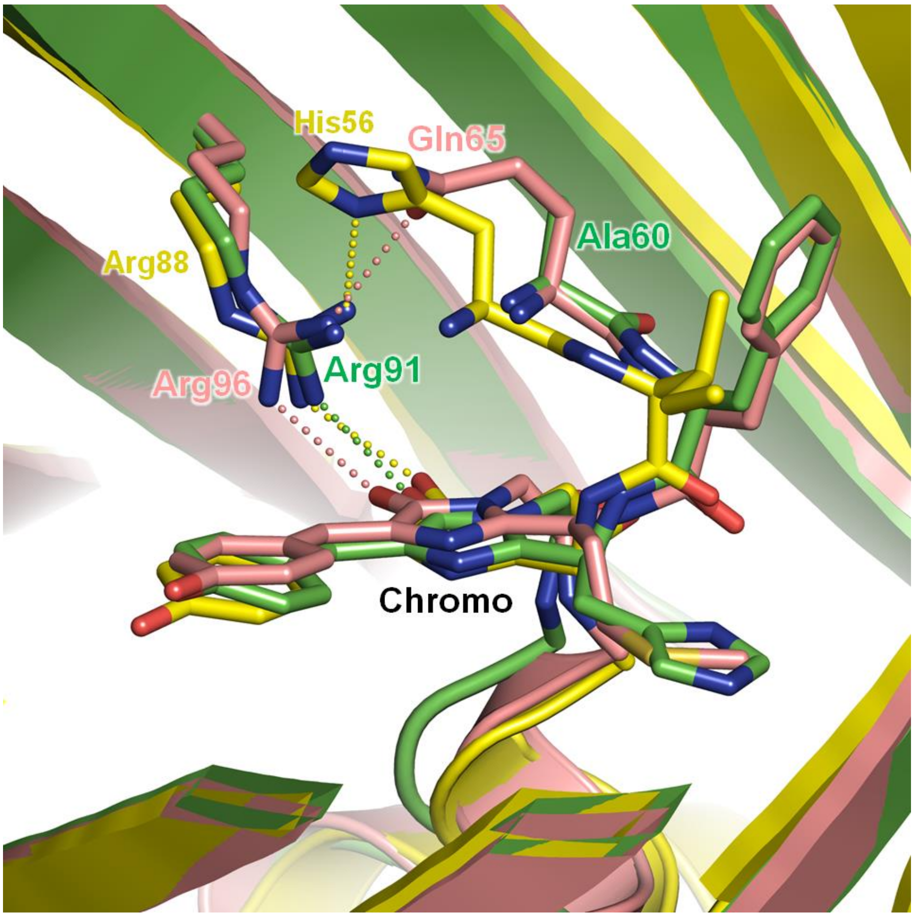


*Figure S4. Structure alignment of mEos4b, mScarlet3 and mNeonGreen.*

*The microenvironment of the chromophore and particularly positions equivalent to A60 and R91 (mEos4b numbering) are shown as green carbons (mEos4b, PDB: 6GOY), pink carbons (mScarlet3, PDB: 7ZCT) and yellow carbons (mNeonGreen, PDB: 5Y00). H-bonds (between 2.8 and 3.1Å) between the arginine and its partners are represented as dashed lines.*

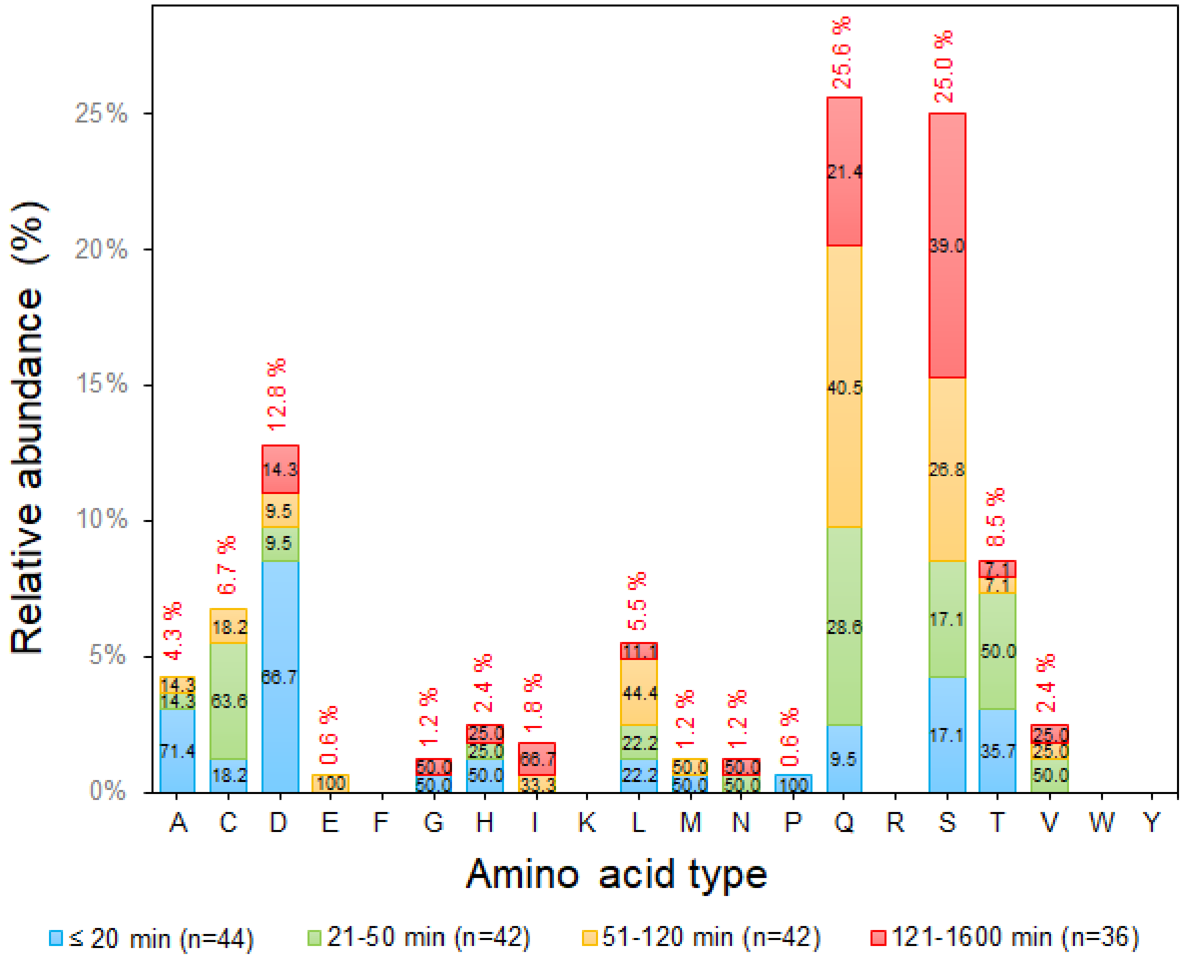


*Figure S5. Survey of the amino acid present at position 60 in function of the maturation time*

*Home-made python scripts were used to parse through FPbase to first filter fluorescent proteins based on a given maturation time cut-off, followed by sequence aligning the proteins to record the amino acid residues present at position 60 (mEos numbering) or equivalent. Results cover all FPs to date with a maturation value reported (164 proteins). Four batches of maturation containing an approximately equal repartition of FPs in each batch have been selected but the last batch gathers a very broad range of significantly slow maturation FPs. Care should be taken in comparing maturations rates of different proteins due to differences in protein classes and wide variety of measurement protocols used.*

*The figure reads as follows: "12.8% of fluorescent proteins have an aspartate (D) at position 60 (mEos4b numbering) and 66.7% of those are maturing in less than 20 min"*

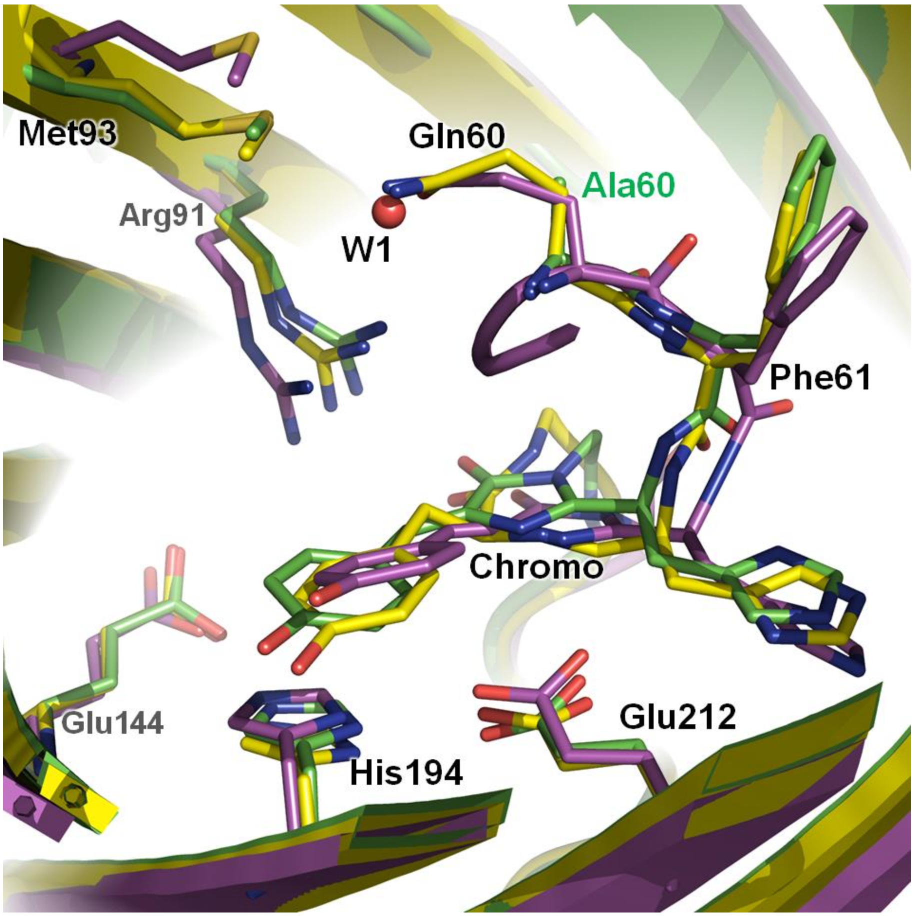


*Figure S6. Structure of mEos4b-L93M and overlaid simulations of its A60Q variant.*

*Simulations of the structures of mEos4b-A60Q-L93M (mEos4Fast1) are represented as yellow carbons (AlphaFold 3 prediction) and purple carbons (Phenix dynamics) and overlaid on the crystallographic structure of mEos4b-L93M. Simulations indicate that Q60 would replace the supplementary water W1 that is present in mEos4b-L93M, providing a partner to both M93 and R91.*

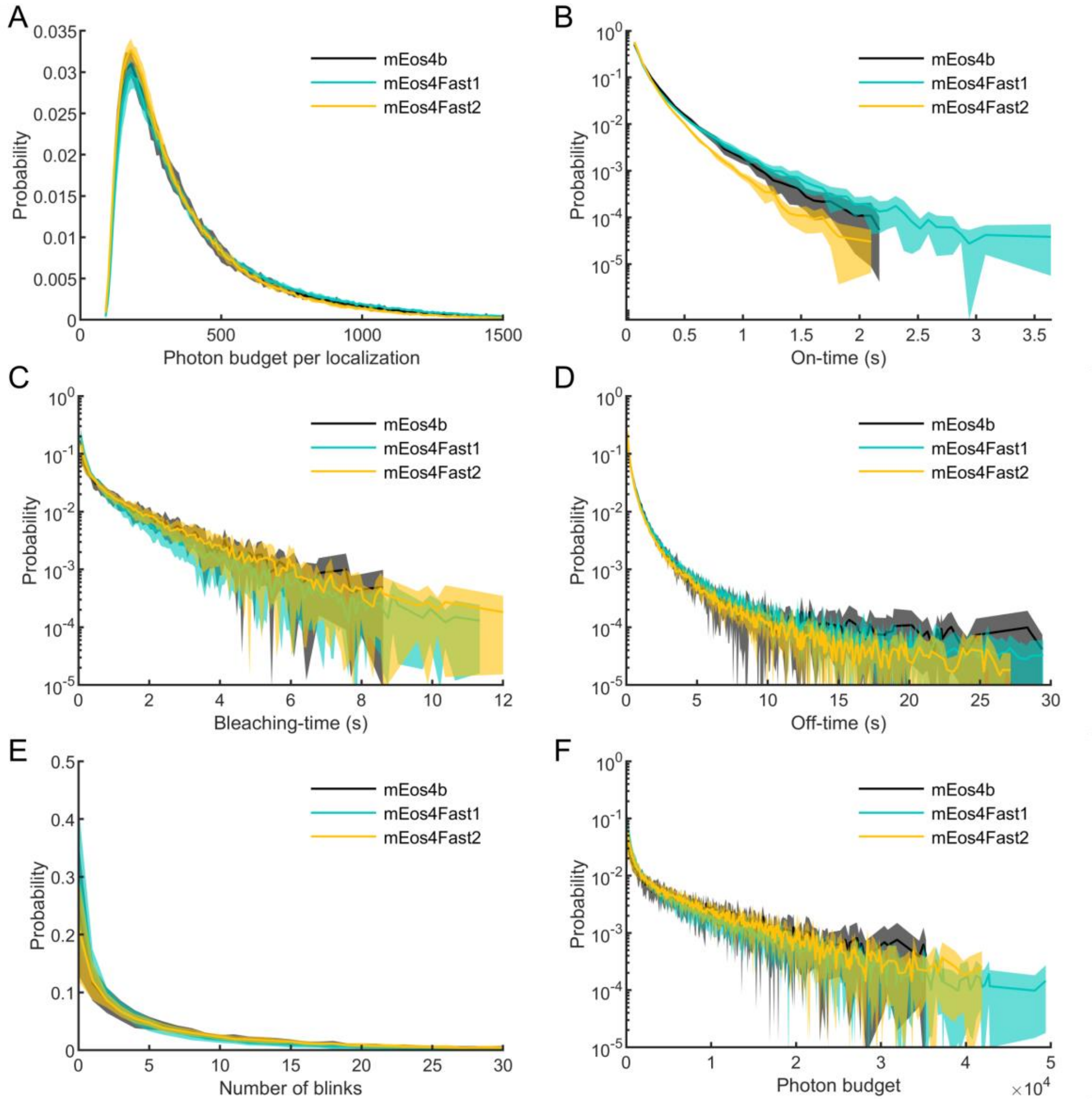


| | Photon budget (s) | On time (s) | Off time (s) | Bleaching time (s) | Number of blinks | Photons per localization |
|---|---|---|---|---|---|---|
| **mEos4b** | 8000.40 ± 2899.14 | 0.17 ± 0.01 | 1.05 ± 0.14 | 1.38 ± 0.51 | 7.03 ± 2.70 | 405.87 ± 15.82 |
| **mEos4Fast1** | 6102.68 ± 2459.54<br>ns | 0.17 ± 0.01<br>ns | 1.01 ± 0.06<br>ns | 1.00 ± 0.38<br>ns | 4.80 ± 2.22<br>ns | 426.21 ± 17.24<br>* |
| **mEos4Fast2** | 7042.82 ± 2714.97<br>ns | 0.14 ± 0.00<br>*** | 0.81 ± 0.04<br>*** | 1.31 ± 0.54<br>ns | 8.27 ± 3.73<br>ns | 379.71 ± 13.53<br>*** |

*Figure S7. Single molecule photophysics*

*PCFPs were immobilized in PAA gel (pH 8) and single molecule imaging was performed using 500 W/cm² 561-nm light (70 ms exposure time) and 1 W/cm² 405-nm light (8.2 ms exposure time). Localizations were clustered to reconstruct fluorescent time traces belonging to single molecules. From these fluorescent time traces, the mean number of photons per localization (A), off-times (B), on-times (C), number of blinks (D), bleaching-times (E) and total photon budget (F) were extracted and are reported in the table. ANOVA was used to compare mEos4Fast1 and mEos4Fast2 against mEos4b (ns = not significant, * p<0.05, *** p<0.0005)*

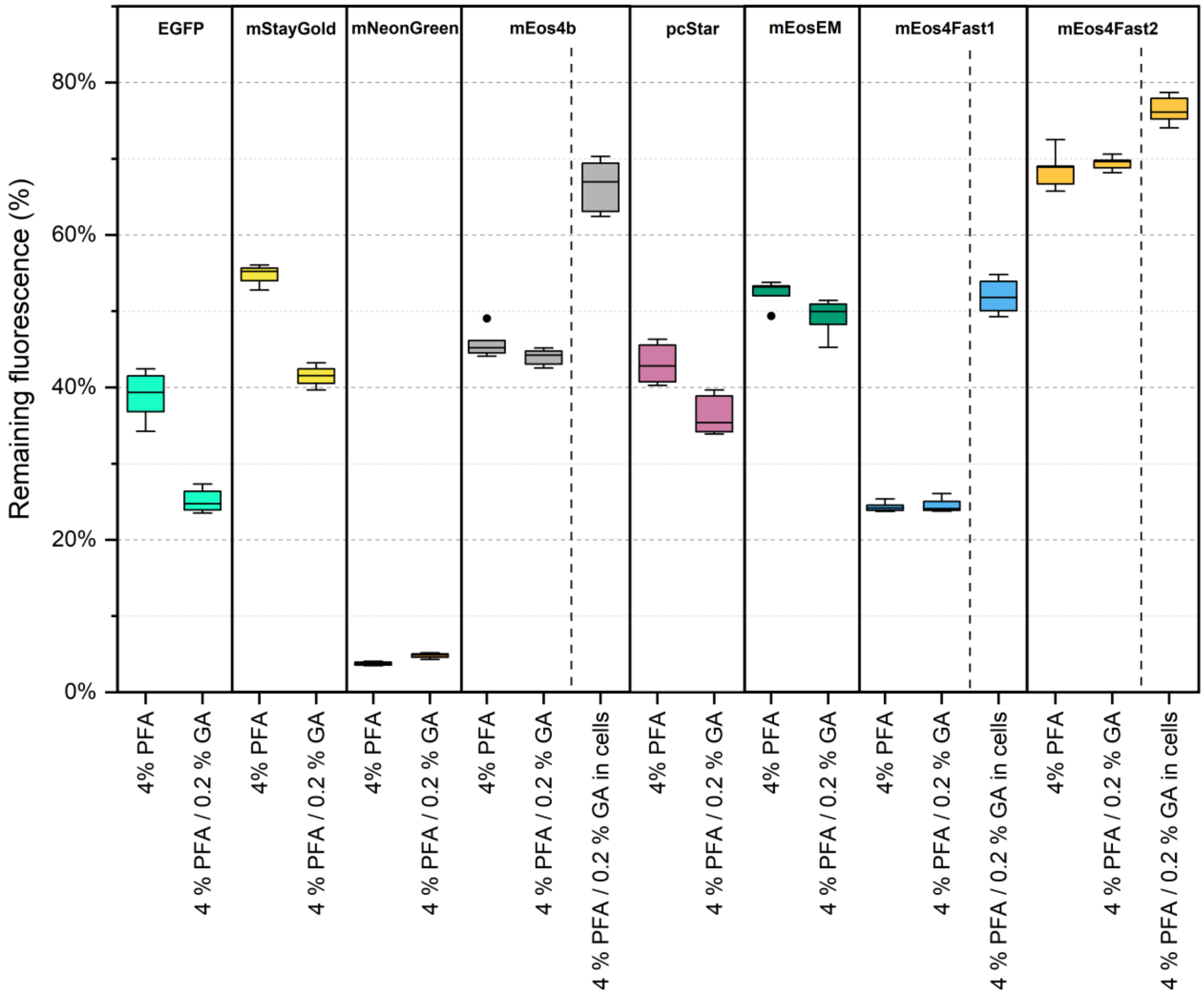


*Figure S8. Resistance to fixation reagents*

*Residual fluorescence levels of several fluorescent proteins in their purified form were assessed following 30 minutes of fixation at 37°C with either 4% paraformaldehyde (PFA) or a combination of 4% PFA and 0.2% glutaraldehyde (GA). The proteins tested include the green FPs EGFP, mStayGold, mNeonGreen as controls and the PCFPs mEos4b, pcStar, mEosEM, mEos4Fast1 and mEos4Fast2 in their green form. The boxplots depict the percentage of remaining fluorescence relative to initial levels, with each protein represented by a different color. Boxes show the interquartile range (IQR), with the horizontal line inside each box indicating the median fluorescence. The whiskers extend to 1.5 times the IQR and points outside this range are considered outliers. n = 6 except for mStayGold and mNeonGreen (n=4). For mEos4b, mEos4Fast1 and mEos4Fast2, fluorescence measurements (median fluorescence intensity) were also performed on U2OS cells 24h post-transfection by flow cytometry (n=6).*

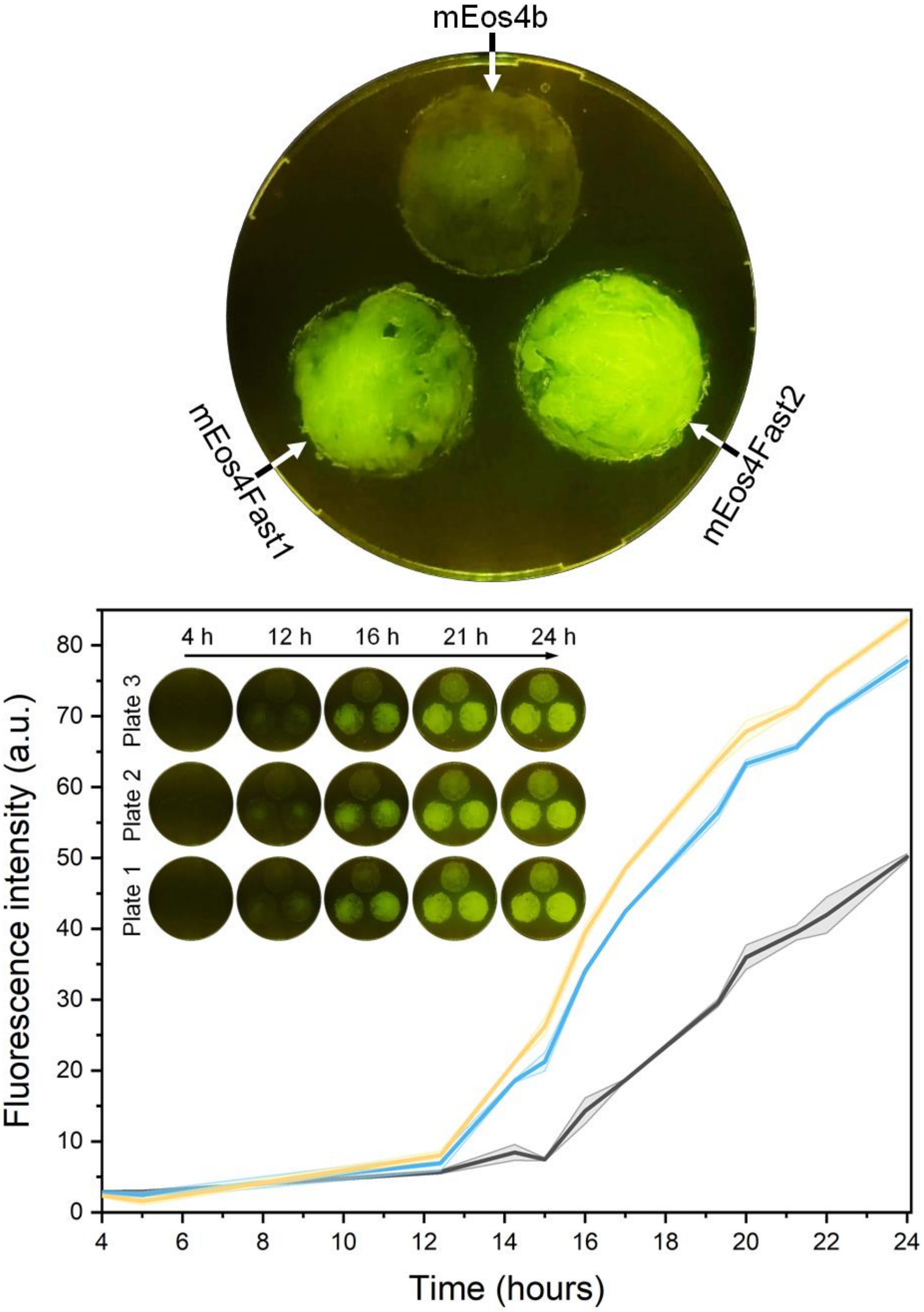


*Figure S9. Fluorescence induction kinetics in E. coli expressing mEos4b, mEos4Fast1 and mEos4Fast2.*

*Top panel: Representative photograph of a Petri dish with E. coli colonies expressing mEos4b, mEos4Fast1 and mEos4Fast2, taken under a blue light transilluminator. Bottom panel: Kinetic curves showing the increase in fluorescence over time for each protein. Data points represent the fluorescence intensity at various time intervals from 4 to 24 hours post-induction with 10 mM IPTG. Inset shows selected time-lapse images of the Petri dishes at specific time points within the 24-hour period. The images illustrate the progressive increase in fluorescence for each protein, highlighting the differences in fluorescence induction and intensity among proteins.*

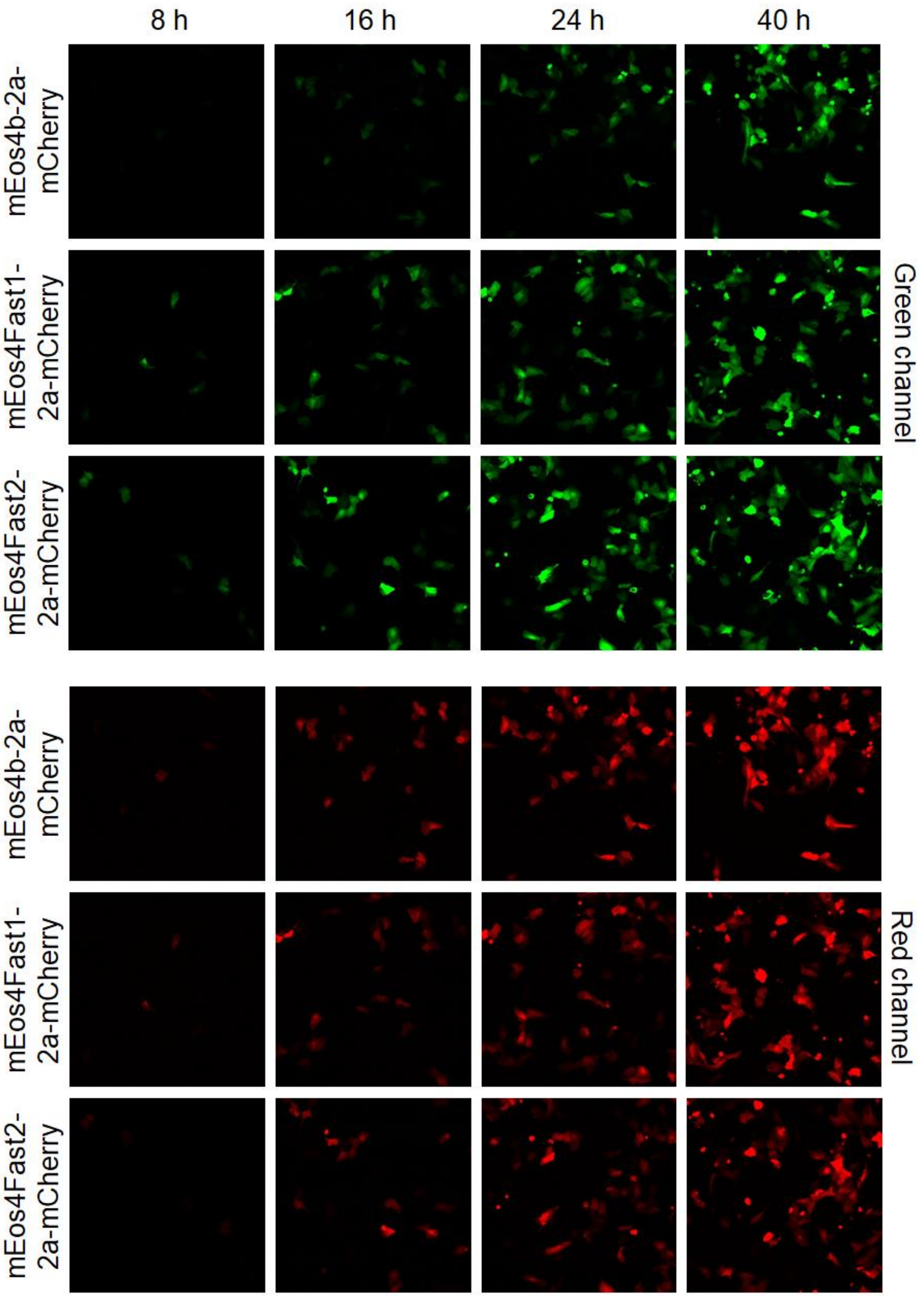


*Figure S10. Maturation of mEos variants in mammalian cells. Evolution of fluorescence signal of U2OS cells transfected with the bicistronic constructs mEos4b-2a-mCherry, mEos4Fast1-2a-mCherry or mEos4Fast2-2a-mCherry. mCherry served as a reference for normalization to account for a difference in transfection and expression levels among samples. Cells were growing in the incubation chamber of the spinning disk microscope (37°C and 5% $CO_2$) and imaged starting 4 h after transfection, for a total duration of 44 h, with a time interval of 30 min. Four represented time points correspond to those of the flow cytometry experiment.*

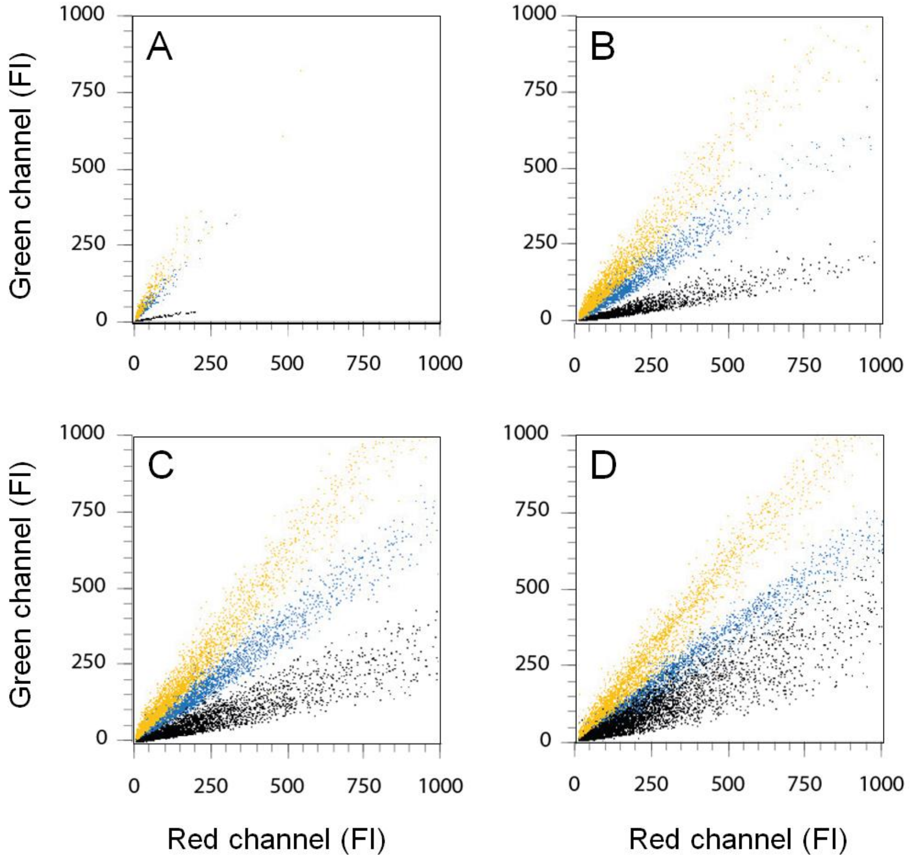


*Figure S11. Flow cytometry 8 hours (A), 16 hours (B), 24 hours (C) and 40 hours (D) post-transfection. Following transfection, cells were harvested and subjected to flow cytometry analysis. Dot plots of mEos4 (green) versus mCherry (red) fluorescence are shown at the indicated times for cells transiently expressing the bicistronic constructs mEos4b-2A-mCherry (black), mEos4Fast1-2A-mCherry (blue), or mEos4Fast2-2A-mCherry (yellow). Fluorescence signals are expressed as arbitrary fluorescence intensity values. A total of 10,000 cells were analysed per condition. FI: Fluorescence Intensity*

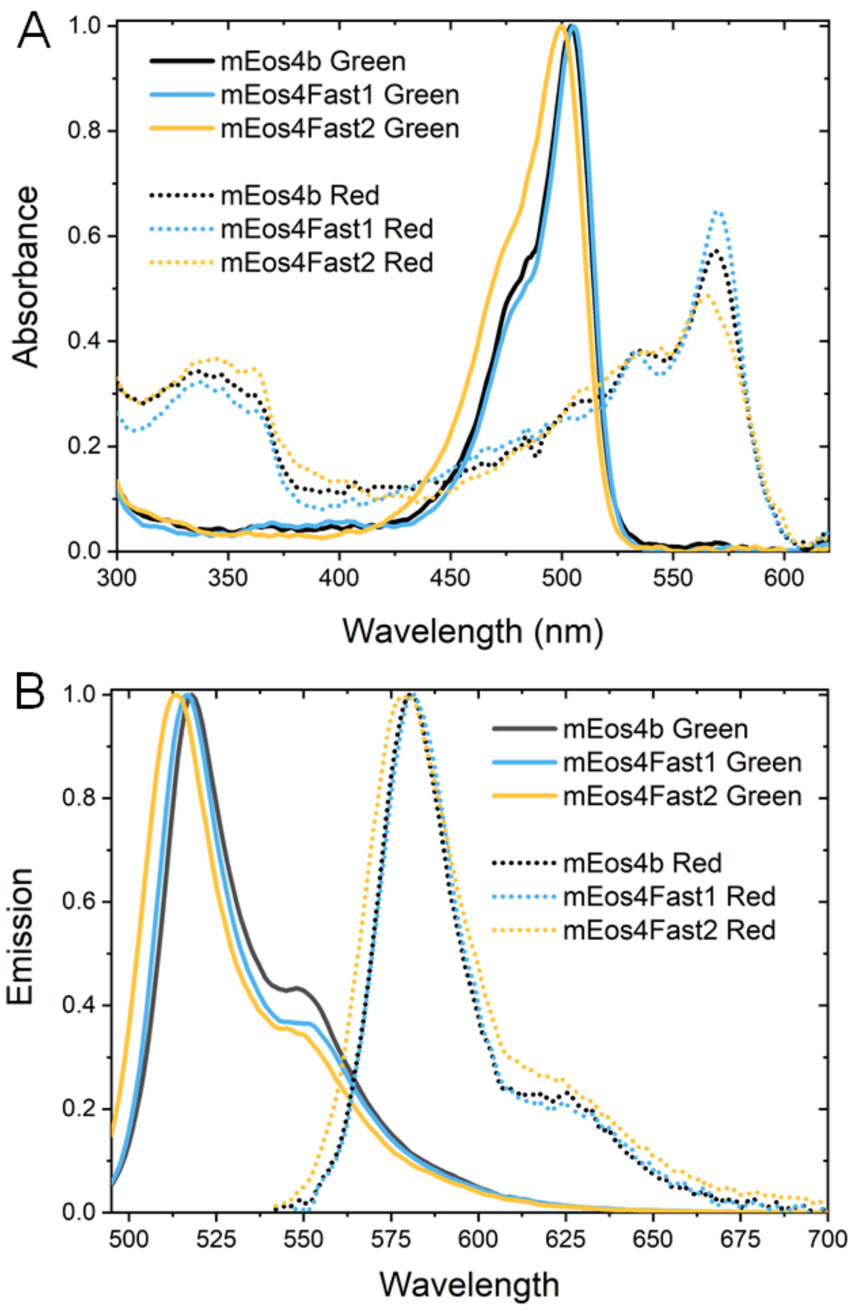


*Figure S12. Absorption (A) and emission (B) spectra of mEos4b, mEos4Fast1 and mEos4Fast2 in their green (plain lines) and red forms (dotted lines).*

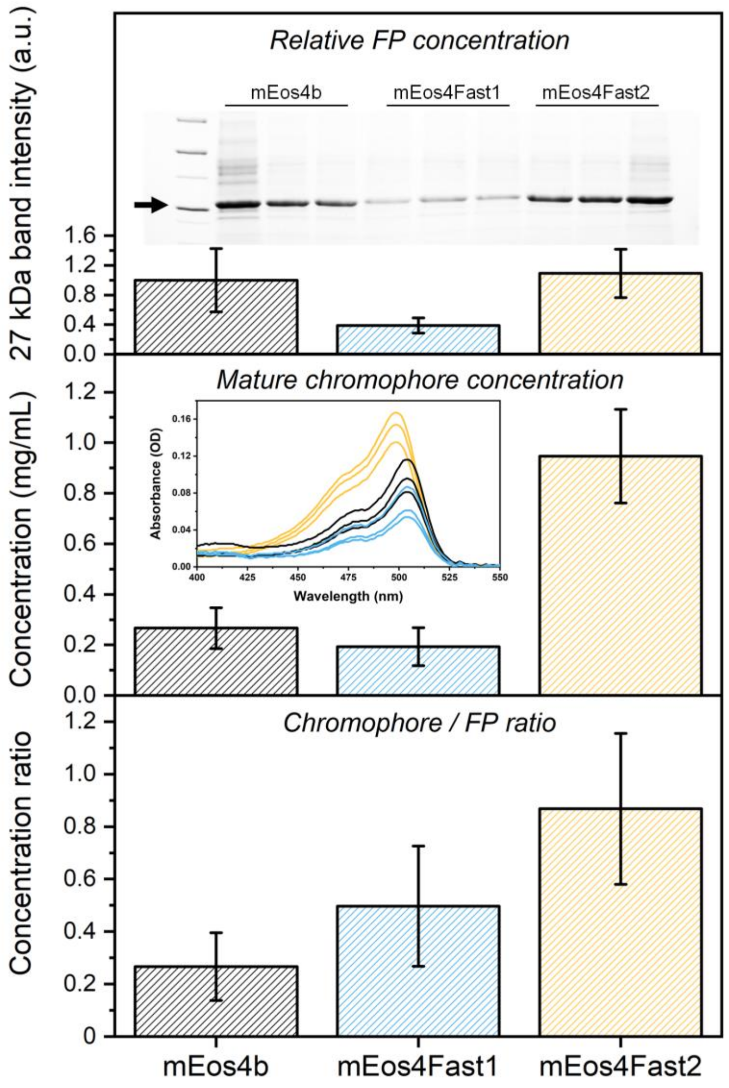


*Figure S13. Protein expression and chromophore maturation in E. coli.*

*Top panel: Integrated 27-kDa band intensity from SDS-PAGE (4-15% gradient in reducing condition) analysis with gel image as inset and showing a lower protein yield for mEos4Fast1. Middle panel: Chromophore concentration measurements with UV-visible absorption spectra as inset and showing a higher yield of chromophores for mEos4Fast2. Bottom panel: Ratios of chromophore concentration to protein concentration, indicating chromophore maturation efficiency. All measurements were made in triplicate. Standard deviations for each measurement is represented by error bars.*

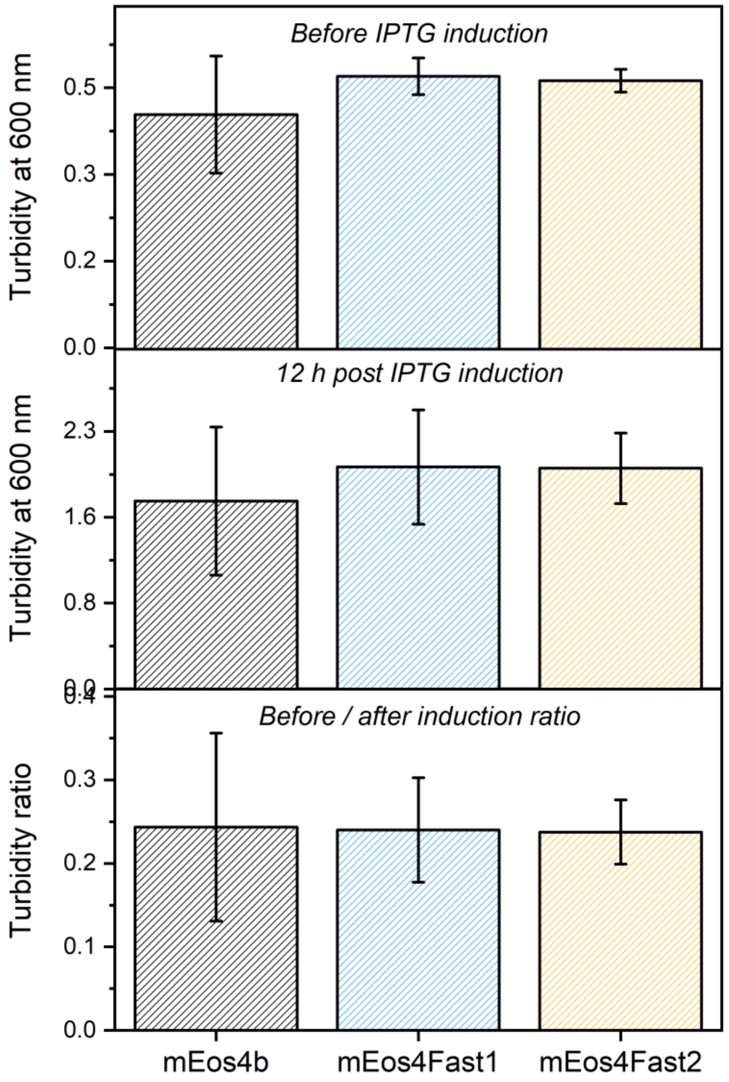


*Figure S14. E. coli turbidity measurements at 600 nm before and after IPTG induction.*

*Top panel: Turbidity (OD600) of cultures before IPTG induction for mEos4b, mEos4Fast1 and mEos4Fast2. Middle panel: Turbidity 12 hours after IPTG induction. Bottom panel: Ratios of turbidity (post-induction/pre-induction) for each protein. The ratios were identical across all three proteins, indicating comparable bacterial growth and expression conditions. Standard deviations for each measurement is represented by error bars.*

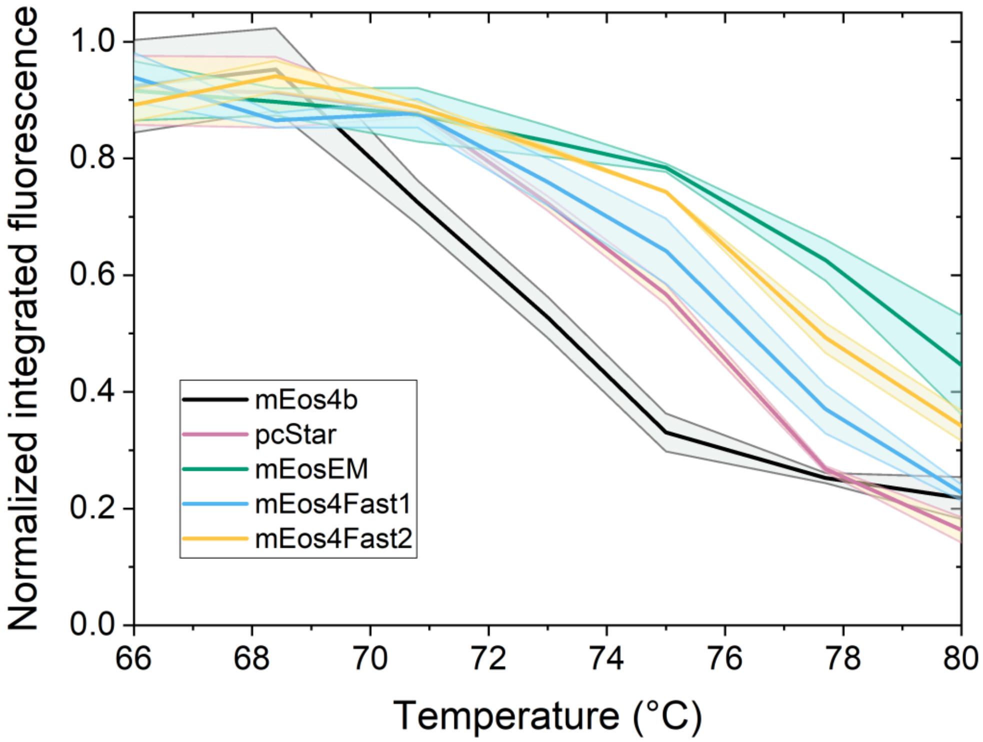


*Figure S15. Thermal stability assay of PCFPs.*

*Residual fluorescence of purified mEos4b, pcStar, mEosEM, mEos4Fast1 and mEos4Fast2 incubated at temperatures ranging from 66°C to 80°C for 30 minutes. Fluorescence was normalised to samples incubated at room temperature. Measurements were performed in triplicate using a multi-well plate reader. mEos4Fast2 demonstrated the second highest thermal resistance, close to mEosEM, indicating substantial stability.*

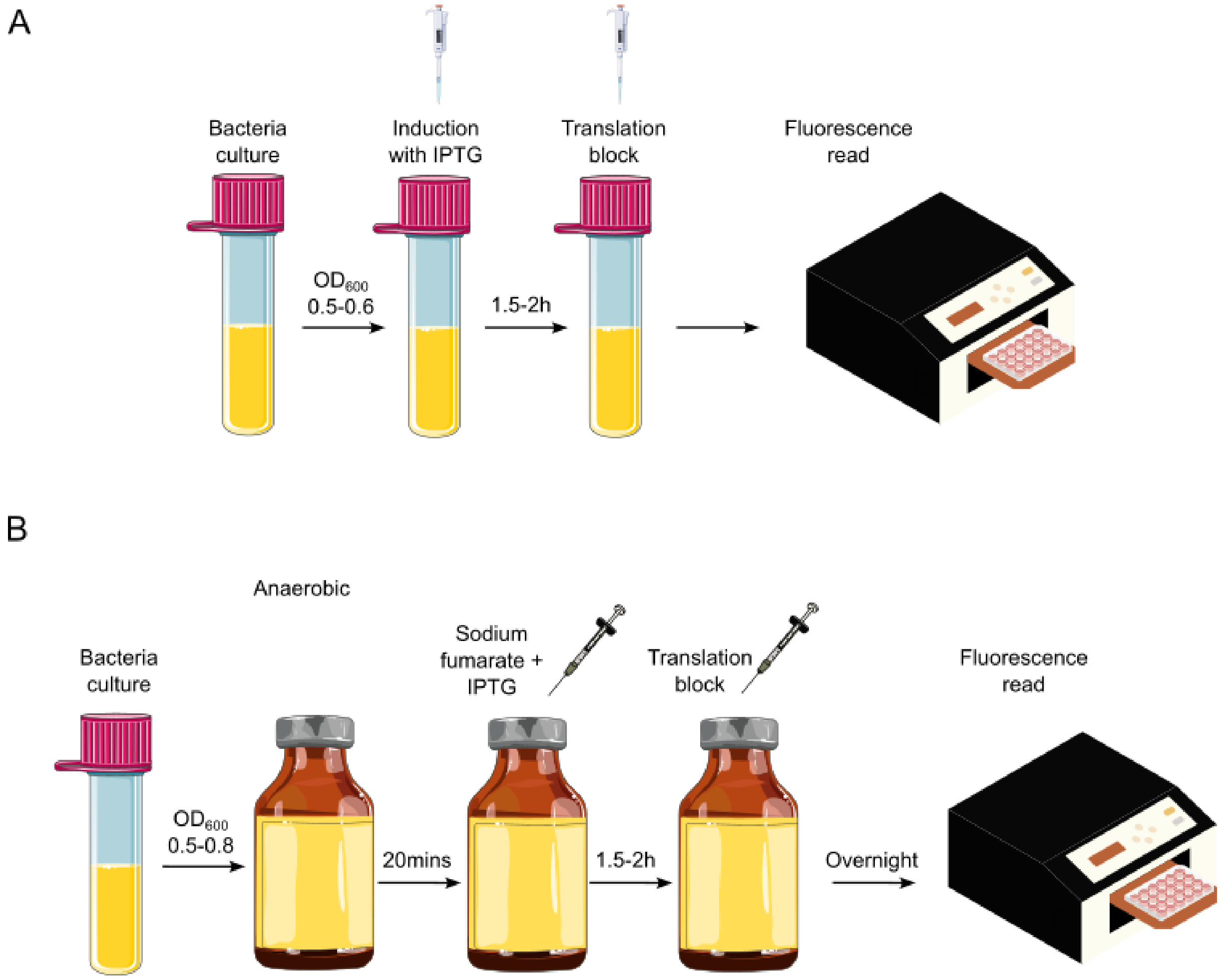


*Figure S16. Graphical illustration of the FP maturation assay performed in A) aerobic condition and B) anaerobic condition. Icons used in this illustration are adapted from https://bioicons.com/*

*Table S1. Comparison of apparent maturation rates in mEos4b and its 60/93 mutants*

| **Protein** | **τ aerobic (min)** | **Protein** | **τ aerobic (min)** |
|---|---|---|---|
| mEos4b | 1860 ± 48 | mEos4b-L93D | NA |
| mEos4b-A60H | NA | mEos4b-L93E | 346 ± 60 |
| mEos4b-A60K | NA | mEos4b-L93M | 81 ± 11 |
| mEos4b-A60P | NA | mEos4b-L93N | 440 ± 54 |
| mEos4b-A60Q | 209 ± 13 | mEos4b-L93Q | 179 ± 10 |
| mEos4b-A60S | 960 ± 80 | mEos4b-A60Q-L93M (mEos4Fast1) | 64 ± 5 |
| mEos4b-A60T | 2133 ± 394 | | |

NA: Not applicable, the considered variants did not mature

*Table S2. List of primers used to generate mutants of the mEos fluorescent proteins presented in this study*

| Mutation | Forward primer | Reverse primer |
|---|---|---|
| V69A, V69T | 5'-CAGGGTATTCRCTAAATATCCAG-3' | 5'-TTGCCGTAATGGAATG-3' |
| I157V | 5'-GACGGGTGATGTTGAGATGGC-3' | 5'-AGCACTCCATCACGCACA-3' |
| L93D, L93E, L93K, L93N, L93Q | 5'-GGAACGAAGCVANACTTTCGAAGACG-3' | 5'-CACGAATACCCCTTAGGAAAC-3' |
| L93M | 5'-GGAACGAAGCATGACTTTCGAAG-3' | |
| A60D | 5'-CCTGACCACTGACTTCCATTACGGCAAC-3' | 5'-ATATCAAAGGCAAAAGGC-3' |
| A60H, A60K, A60Q | 5'-CCTGACCACTMASTTCCATTACGGCAAC-3' | |
| A60P | 5'-CCTGACCACTCCATTCCATTACGGCAAC-3' | |
| A60S, A60T | 5'-CCTGACCACTWCATTCCATTACGGCAAC-3' | |
| T59Q | 5'-TATCCTGACCCARGCATTCCATTACG-3' | 5'-TCAAAGGCAAAAGGC-3' |
| T59V | 5'-TATCCTGACCGTNGCATTCCATTACG-3' | |

*Table S3. Data collection and refinement statistics of mEos4b-L93M*

| | |
|---|---|
| **PDB code** | 9GVR |
| **Beamline** | ID30-A3/MASSIF3 (ESRF) |
| **Wavelength** | 0.9677 |
| **Resolution range** | 36.69 - 1.864 (1.93 - 1.864) |
| **Space group** | P 21 21 21 |
| **Unit cell** | 39.28 57.30 102.75 90 90 90 |
| **Total reflections** | 167905 (17129) |
| **Unique reflections** | 20036 (1913) |
| **Multiplicity** | 8.4 (8.8) |
| **Completeness (%)** | 99.69 (97.80) |
| **Mean I/sigma(I)** | 8.92 (2.46) |
| **Wilson B-factor** | 11.59 |
| **R-merge** | 0.2225 (1.242) |
| **R-meas** | 0.2373 (1.32) |
| **R-pim** | 0.08116 (0.442) |
| **CC1/2** | 0.993 (0.718) |
| **CC*** | 0.998 (0.914) |
| **Reflections used in refinement** | 19991 (1913) |
| **Reflections used for R-free** | 2000 (191) |
| **R-work** | 0.1681 (0.2172) |
| **R-free** | 0.2071 (0.3133) |
| **CC(work)** | 0.954 (0.784) |
| **CC(free)** | 0.944 (0.652) |
| **Number of non-hydrogen atoms** | 2099 |
| **macromolecules** | 1807 |
| **ligands** | 52 |
| **solvent** | 240 |
| **Protein residues** | 217 |
| **RMS(bonds)** | 0.004 |
| **RMS(angles)** | 0.78 |
| **Ramachandran favored (%)** | 99.06 |
| **Ramachandran allowed (%)** | 0.94 |
| **Ramachandran outliers (%)** | 0.00 |
| **Rotamer outliers (%)** | 1.55 |
| **Clashscore** | 3.01 |
| **Average B-factor** | 14.63 |
| **macromolecules** | 12.86 |
| **ligands** | 24.10 |
| **solvent** | 25.95 |

Statistics for the highest-resolution shell are shown in parentheses.
